# Improved Detection of Drug-Induced Liver Injury by Integrating Predicted *in vivo* and *in vitro* Data

**DOI:** 10.1101/2024.01.10.575128

**Authors:** Srijit Seal, Dominic P. Williams, Layla Hosseini-Gerami, Manas Mahale, Anne E. Carpenter, Ola Spjuth, Andreas Bender

## Abstract

Drug-induced liver injury (DILI) has been significant challenge in drug discovery, often leading to clinical trial failures and necessitating drug withdrawals. The existing suite of in vitro proxy-DILI assays is generally effective at identifying compounds with hepatotoxicity. However, there is considerable interest in enhancing in silico prediction of DILI because it allows for the evaluation of large sets of compounds more quickly and cost-effectively, particularly in the early stages of projects. In this study, we aim to study ML models for DILI prediction that first predicts nine proxy-DILI labels and then uses them as features in addition to chemical structural features to predict DILI. The features include *in vitro* (e.g., mitochondrial toxicity, bile salt export pump inhibition) data, *in vivo* (e.g., preclinical rat hepatotoxicity studies) data, pharmacokinetic parameters of maximum concentration, structural fingerprints, and physicochemical parameters. We trained DILI-prediction models on 888 compounds from the DILIst dataset and tested on a held-out external test set of 223 compounds from DILIst dataset. The best model, DILIPredictor, attained an AUC-ROC of 0.79. This model enabled the detection of top 25 toxic compounds compared to models using only structural features (2.68 LR+ score). Using feature interpretation from DILIPredictor, we were able to identify the chemical substructures causing DILI as well as differentiate cases DILI is caused by compounds in animals but not in humans. For example, DILIPredictor correctly recognized 2-butoxyethanol as non-toxic in humans despite its hepatotoxicity in mice models. Overall, the DILIPredictor model improves the detection of compounds causing DILI with an improved differentiation between animal and human sensitivity as well as the potential for mechanism evaluation. DILIPredictor is publicly available at https://broad.io/DILIPredictor for use *via* web interface and with all code available for download and local implementation via https://pypi.org/project/dilipred/.

**GRAPHICAL ABSTRACT:** 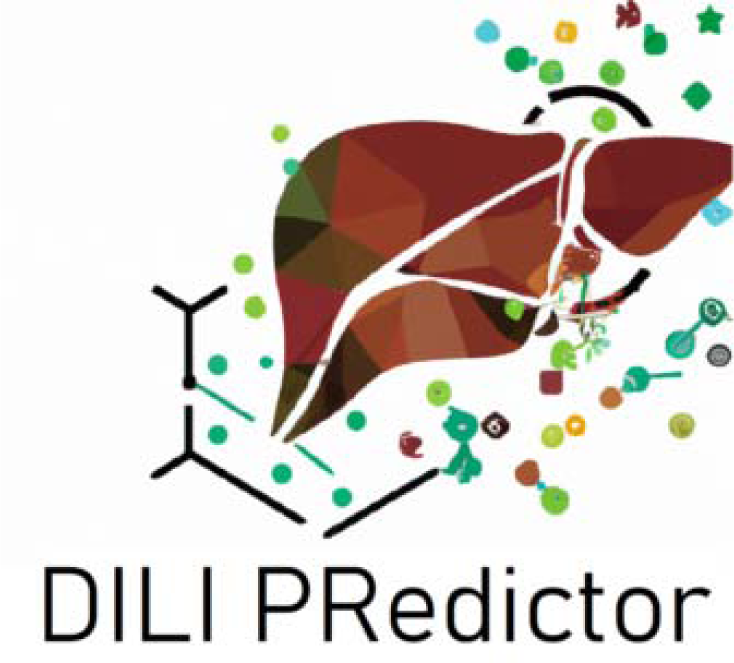

## INTRODUCTION

The liver is a major organ of drug metabolism in the human body and thus it is vulnerable to not just drugs but also their reactive metabolites.^1,2^ Drug-induced liver injury (DILI) has been a leading cause of acute liver failure^3^ (causing over 50% of such cases^4^), accounting for a significant proportion of drug-related adverse events. DILI is detectable generally in phase III clinical trials and a leading cause of post-market drug withdrawals.^5^ Two common types of DILI are intrinsic and idiosyncratic.^6^ While intrinsic DILI is generally dose-dependent and predictable, idiosyncratic DILI is rare, generally not dependent on dosage, typically unpredictable undetectable in early drug development using standard preclinical models, with a variable onset time and several phenotypes.

The mechanisms underlying DILI are multifactorial^7^ and not completely understood. These include cellular toxicities such as mitochondrial impairment^8^, inhibition of biliary efflux^9^, oxidative stress^10^, and more. Additionally, DILI can be influenced by dose variations, pharmacokinetics (PK), and biological variations, such as variations in cytochrome P450 (CYP) expression^11^. Today, there are several established *in vitro* assays and *in vivo* experiments (proxy-DILI data) that are relatively good at detecting DILI risk.^12^ The current battery of *in vitro* proxy-DILI assays is generally effective at detecting hepatotoxic compounds. However, most human-relevant in vitro models for DILI risk assessments can still fail to accurately predict patient safety at therapeutic doses due to differing toxicity mechanisms at varying concentrations. At the time of writing this research article (February - June 2024), at least three reports show clinical trials in phase 1/2 suffered setbacks or ended due to unexpected liver damage, liver function abnormalities that was not predicted by early in vitro models.^13,14,15^ This underscores the critical need for the development of more accurate and reliable in vitro systems that can better simulate human liver responses to drug exposure, thereby improving the safety assessment and reducing the risk of late-stage clinical failures. Thus, there is significant interest in improving *in silico* DILI prediction due to its ability to assess large numbers of compounds more quickly and cost-efficiently, especially in the early stages of drug discovery.^16^

In the drug discovery pipeline, hepatotoxicity assessment encompasses a variety of *in vitro* and *in vivo* experimental models as well as *in silico* models. Several *in vitro* models for liver toxicity testing employ proxy endpoints (hepatotoxicity assays) with liver slices and cell lines such as primary animal and human hepatocytes^17^ or even three-dimensional systems with the dynamic flow for the primary cell and/or stem cell cultures.^18^ However, the ideal hepatocyte-like cell model system depends on the evaluation of particular cellular functions given there are substantial differences among various human liver-derived single-cell culture models as previously explored in the context of drug disposition, bioactivation, and detoxification.^19^ The agreement between *in vitro* data and human *in vivo* data is also low.^20^ For example, methapyrilene is known to cause changes to the level of iron metabolism in the human hepatic HepaRG cell line^21^ and oxidative stress, and mitochondrial dysfunction in rats^22^ but has not been reported to cause hepatotoxicity in humans^23,24^. On the other hand, *in vivo* animal models also have low concordance as shown by recent studies using the eTOX database where organ toxicities were rarely concordant between species.^25^ The concordance between animal and human data for liver toxicity, specifically, is often low (with some studies indicating rates as low as 40%^26^ and others in the range of 39−44%^27^) which makes extrapolating safety assessments from animals to humans a challenging endeavour.^28,29^ For example, 2-butoxyethanol causes hepatic toxicity in mice *via* an oxidative stress mechanism but not in humans given humans have higher levels of liver vitamin E (and a high resistance to iron accumulation) compared to mice.^30^ Overall, this leads to a greater need for improved DILI prediction, especially when translating knowledge from preclinical stage and animal studies to human clinical studies.^16^

DILIst^31^ and DILIrank^32^ are lists of compounds that have been classified as inducing DILI or not and were developed from FDA-approved drug labels. Binary classification from labelling documents is challenging and this is evident in the fact that many DILIrank compounds are labelled ambiguous although the DILI for some of these compounds has been reported in literature. For *in silico* models, these ambiguous compounds are generally removed. Machine learning models are being increasingly used to model biological systems and identify complex patterns in datasets.^33^ Generally, *in silico* models rely on identifying chemical structural alerts^34^ or use a range of chemical or physicochemical features. Ye et al. employed Random Forest algorithms and Morgan fingerprints for DILI prediction, achieving an AUC of 0.75 with random splitting (70% training, 30% testing).^35^ Liu et al. utilized Support Vector Machines and obtained a 76% balanced accuracy on an external test set using Morgan Fingerprints; however, their predicted protein target descriptors provided less accurate predictions (balanced accuracy of 59%) but offered better interpretability.^36^ Mora et al. employed QuBiLS-MAS 0–2.5D molecular descriptors to predict DILI (labels from various sources) on an external test set comprising 554 compounds, achieving a 77% balanced accuracy.^37^ Predicting organ-level toxicity solely based on chemical structure is challenging and the use of biological data helps improve toxicity prediction.^38,39^ More recently, predicted off-target effects and experimental P450-inhibitory activity have also been considered to improve DILI prediction.^40,41^ Moving away from binary predictions, Aleo et al. developed the hepatic risk matrix (HRM) to assess the potential for human drug-induced liver injury (DILI) among lead clinical and back-up drug candidates. By integrating physicochemical properties and common toxicity mechanisms, the HRM stratifies drug candidates based on safety margins relative to clinical C_max,total_. This study identified 70-80% most-DILI-concern drugs and effectively differentiated successful from unsuccessful drug candidates for liver safety.^42^ Chavan et al. integrated high-content imaging features with chemical features for DILI label prediction, resulting in a 0.74 AUC.^43^ Previously this, the authors of this work explored this in the case of mitochondrial toxicity^44^ (which at high doses is one of the mechanisms known to cause DILI), cytotoxicity^45^, as well as cardiotoxicity^46^.

In this study, we significantly extended the use of different data sources to several *in vivo* and *in vitro* data types in developing the DILIPredictor model presented here. We identified liver injury endpoints such as human hepatotoxicity^47^, preclinical hepatotoxicity and animal hepatotoxicity^47,48,49^ and DILI datasets compiled by various studies^37,50^ (Table 1) These datasets provide the *in vivo* labels for DILI for different species at various stages of the drug discovery pipeline, from pre-clinical to post-market withdrawals. We identified three *in vitro* assays that could be indicative of liver toxicity and with public data^7^: mitochondrial toxicity^51^, bile salt export pump inhibition (BSEP)^52^ and the formation of reactive metabolites^53^. Mitochondria accounts for 13-20% of the liver, and mitochondrial dysfunction can impact ATP synthesis, increase ROS generation and trigger liver injury.^54^ The majority of the mitochondrial toxicity data in Hemmerich et al. originates from a Tox21 assay assessing mitochondrial membrane depolarization in HepG2 cells (which provides a distinct perspective compared to *in vitro* data derived from primary hepatocytes) thereby introducing additional biological information. When BSEP function is inhibited, bile salts accumulate within liver cells, causing hepatocyte injury and a risk of liver failure.^55^ Metabolic processes can form reactive metabolites that bind covalently to hepatic proteins, altering their function and leading to damage in liver tissues.^56^ We also included PK parameters, which have been predicted before using machine learning models based on chemical structures.^57^ Overall, in this study, we hypothesized that these proxy-DILI labels along with chemical structure and physicochemical parameters would lead to imrpoved predictivity in identifying potential liver injury endpoints while differentiating between sensitivities observed in human and animal proxy-DILI labels, allowing for interpretations of hepatotoxicity data across species. An objective of our study is not just to achieve high overall predictive performance, but to understand how individual in vitro and in vivo proxy endpoints provide predictive value for DILI outcomes in humans. The prediction of individual proxy endpoints is valuable, as it aligns with the specific experiments, thereby forming testable hypotheses. It should be noted that in vivo DILI is not easily testable without significant effort and clinical trials. Hence in this work, a FeatureNet approach was adopted, that is models are trained on predictions from other individual models. This allows us to interpret the importance of the individual predictions to the final prediction with previous studies showing comparable performance to multitask learning.^58^ Furthermore, the nature of our data, characterized by small sizes for in vivo relevant data and sparse matrices, presents additional challenges for implementing multitask learning effectively. Finally, by including *in vitro* proxy-DILI labels, the models developed in this study have the potential for mechanistic evaluation and facilitating a comprehensive understanding of the underlying biochemical and cellular processes associated with drug-induced liver injuries.

**Table 1.**
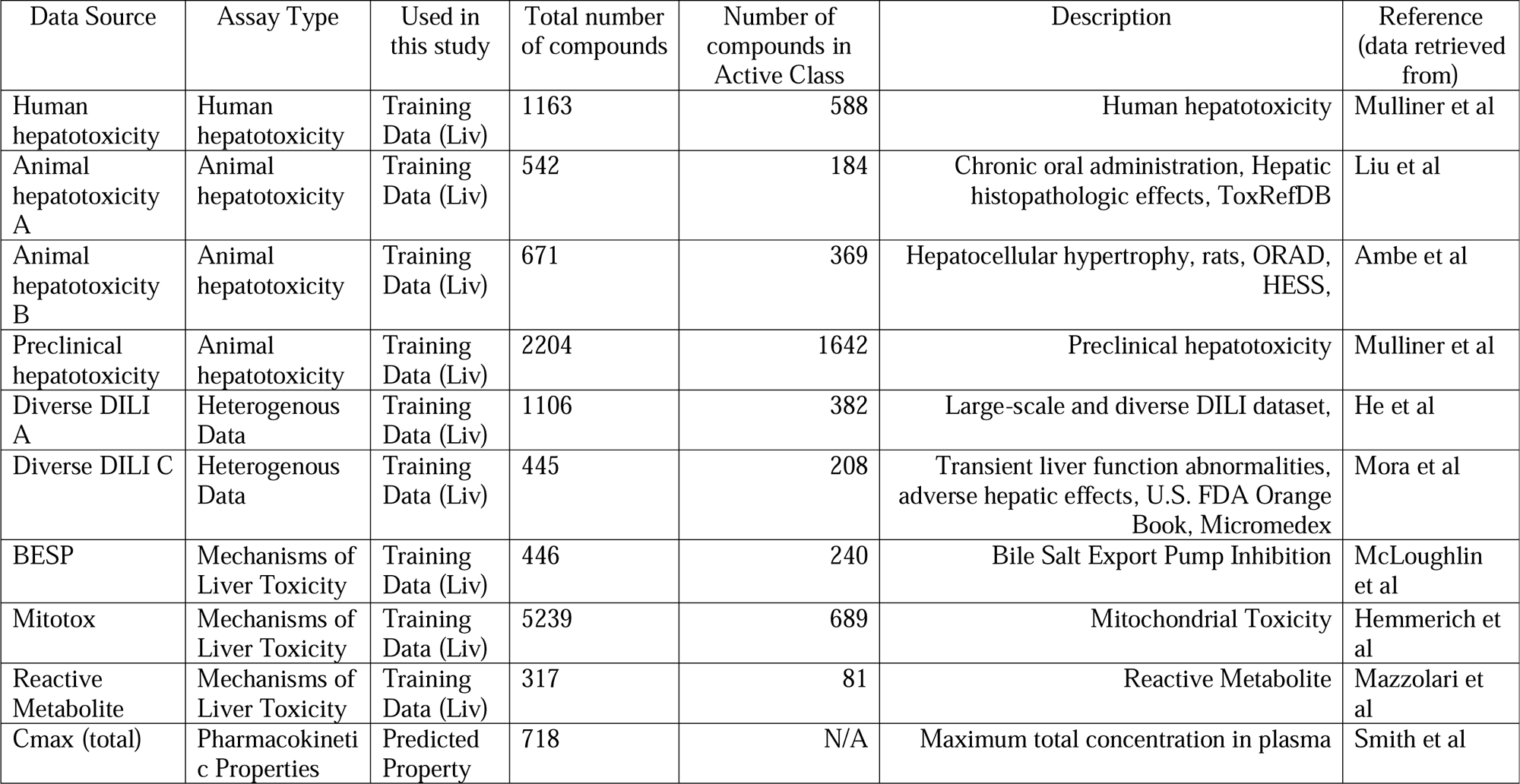
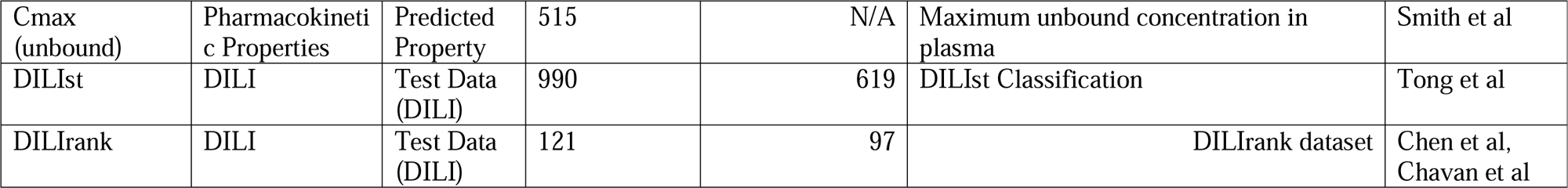
Sources of Liver-safety and Toxicity Data Used in this study.

## MATERIALS AND METHODS

The workflow followed in this study is shown in Figure 1 and described in more detail in the following.

**Figure 1:**
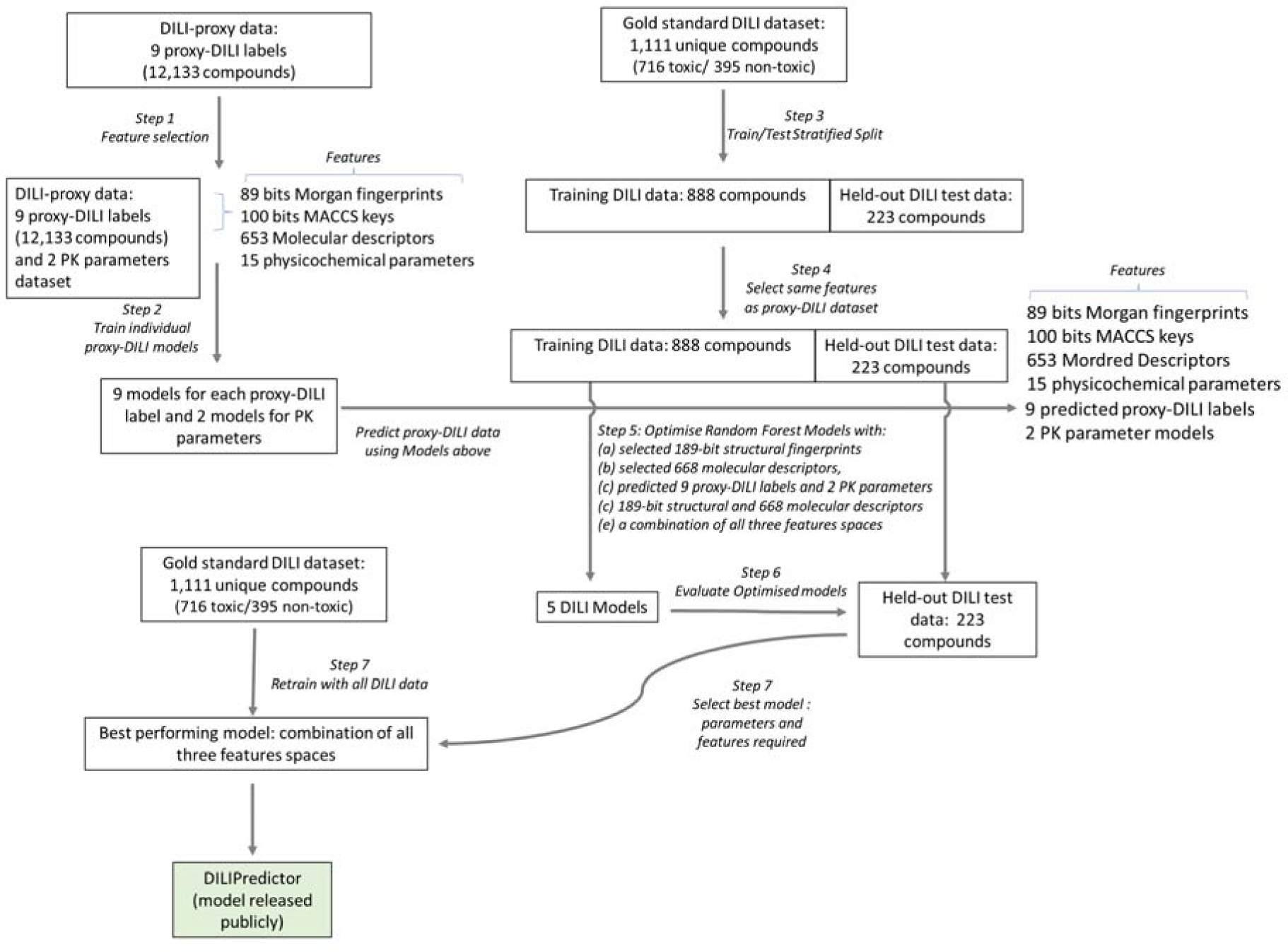
Workflow of the current study. Individual models for 9 *in vivo* and *in vitro* assays in the Proxy-DILI dataset and 2 PK parameters were used to predict these endpoints for compounds in the gold standard DILI dataset. A combination of these predictions along with chemical structure and molecular descriptors were then used to train and evaluate the models on DILI compounds.

### Drug-Induced Liver Toxicity Datasets: DILIst and DILIrank

The human *in vivo* dataset for liver toxicity was collected by combining DILIst^31^ (714 toxic and 440 non-toxic compounds) and DILIrank^32^ (268 toxic and 76 nontoxic compounds from Chavan et al^38^) datasets. The DILIst dataset classifies compounds into two classes based on their potential for causing DILI. The DILIrank dataset was released by the FDA prior to DILIst. This dataset analysed the hepatotoxic descriptions from FDA-approved drugs and assessed causality evidence from literature and classified compounds into four groups: ^v^Most-, ^v^Less, ^v^No-DILI concern and Ambiguous-DILI-concern drugs. For the DILIrank dataset, we retrieved data from Chavan et al^38^. We treated ^v^Most- and ^v^Less as DILI Positive and those labelled with ^v^No-DILI-concern as DILI Negative. Ambiguous-DILI-concern drugs were removed. Together these datasets form the largest drug list with DILI classification to date.

### Proxy-DILI datasets: *in vivo* and *in vitro* assays

The datasets we considered include one proxy-DILI label from studies on human hepatotoxicity^47^, and two proxy-DILI labels from animal hepatotoxicity studies (animal hepatotoxicity A and B, and preclinical hepatotoxicity as detailed in Table 1)^47,48,49^. Animal hepatotoxicity datasets mentioned above consisted of data compiled by the authors from ToxRefDB^59^, ORAD^49^ and HESS^60^ as well as hepatic histopathologic effects. Two diverse DILI datasets contain heterogeneous data collected by other studies^37,50^ (Diverse DILI A and C as detailed in Table 1). These datasets consisted of data from the drugs known to cause transient liver function abnormalities and adverse hepatic effects as well as compounds from the U.S. FDA Orange Book and Micromedex. We included three *in vitro* assays related to proposed or known mechanisms of liver injury, namely, mitochondrial toxicity^51^, bile salt export pump inhibition (BSEP)^52^ and the formation of reactive metabolites^53^ (as detailed in Table 1). The labels for these *in vitro* datasets were the assay hit calls defined by the original studies. Supplementary Table S1 lists the SMILES, chemical names and CASRN for all molecules, where available from the original sources; note that not all original sources provide complete information on chemical names or CASRN. Previous studies indicated that mitochondrial toxicity and BSEP are reasonable predictors for cholestatic and mitochondrial toxins, however, they fail when applied to a wider chemical space for drugs with different mechanisms.^61^ Many assay hits screened from chemical libraries often have unfavourable drug metabolism and pharmacokinetics presenting development challenges.^62^ Thus, we considered pharmacokinetics as two of the proxy-DILI labels and compiled pharmacokinetic parameters of maximum concentration (Cmax) from Smith et al.^63^ This dataset contains maximum unbound concentration in plasma for 515 unique compounds and maximum total concentration in plasma for 718 unique compounds (Supplementary Table S2). Together, as shown in Table 1, we obtained nine *in vivo* and *in vitro* assays (proxy-DILI labels) related to liver injury and two pharmacokinetic parameters.

### Dataset pre-processing and Compound Standardisation

So that the focus of this work remains only on traditional drug like molecules, we dropped compounds if there were no carbon atoms present, if they were disconnected compounds, if they contained contain only metals, or if the molecular weight was above 1500 Daltons. Then, compound SMILES were standardized using RDKit^64^ which included disconnection of metal atoms, normalization, and reionization, isolating the principal molecular fragment, uncharging and charge normalization, standardization to most common isotopic form, removing stereochemical configuration and tautomer standardization. The entire process was iteratively applied up to five times or until the SMILES representation stabilized. If the standardization process did not converge to a single representation, the most frequently occurring SMILES string across the iterations was selected as the standardized form. Finally, the standardized SMILES were protonated to reflect their form at physiological pH of the liver (pH = 7.0) implemented using Dimorphite-DL.^65^ To ensure our dataset did not contain duplicate chemistry, we used the first 14 characters of InChIKeys (known as the “hash layer”) which represents the chemical structure excluding stereochemical and isotopic variations. By comparing these truncated InChIKeys, we identified duplicates. In cases where DILIrank and DILIst contained the same compound with conflicting toxicity labels, we retained the toxic annotation (given there was some evidence of toxicity in either DILISt or DILIrank). Finally, we obtained a dataset of 1,111 unique compounds and associated DILI labels (716 toxic and 395 non-toxic compounds). This dataset is henceforth referred to as the gold standard DILI dataset (Supplementary Table S3).

For the proxy-DILI dataset, in case of any compounds with conflicting toxicity labels within a particular dataset after SMILES standardisation, we retained the compound as toxic/active (hence preferring the evidence of toxicity/activity which is a usual practice in drug discovery) resulting in a dataset of 13,703 compounds. For each of the nine labels besides PK parameters (as detailed in Table 1), if a compound was already present in the gold standard DILI dataset above (compared using the InChIKey hash layer), we removed the compound from the proxy-DILI dataset. This was done to avoid any information leaks in the models developed in this study. Finally, we obtained a dataset of 12,133 compounds in total for nine proxy-DILI labels which are henceforth called the proxy-DILI dataset in this study (Supplementary Table S4).

### Assay Concordance with Experimental Values

To evaluate the concordance of the nine proxy-DILI labels and the gold standard DILI dataset with each other, we used all 13,703 compounds in the proxy-DILI dataset and compared them to the 1,111 compounds in the gold standard DILI dataset. To evaluate concordance, we used Cohen’s kappa (as defined in scikit-learn v1.1.1^66^) to measure the level of agreement between activity values for each pair of labels which were present in the dataset.

### Exploring the Physicochemical Space

Physicochemical space was explored using six characteristic physicochemical descriptors of molecular weight, TPSA, number of rotatable bonds, number of H donors, number of H acceptors and log P, (as implemented in RDKit^64^ v.2022.09.5). We used a t-distributed stochastic neighbour embedding (t-SNE from scikit-learn v1.1.1^66^) to obtain a map of the physicochemical space for all compounds in the gold standard DILI dataset and proxy-DILI dataset with a high explained variance (PCA: 85.15% using two components).

### Structural fingerprints, Mordred, and Physicochemical Descriptors

We used Morgan Fingerprints^67^ of radius 2 and 2048 bits and 166-bit MACCS Keys^68^, as implemented in RDKit^64^ (v2022.09.5), as structural features for all compounds in the DILI dataset and proxy-DILI dataset. This resulted in 2,214-bit vector structural fingerprints.

We used molecular descriptors (as implemented in the Mordred^69^ python package) and physicochemical properties (such as topological polar surface area TPSA, partition coefficient log P etc. as implemented in RDKit^64^ v2022.09.5) for all compounds in the gold standard DILI dataset and proxy-DILI dataset. We dropped descriptors with missing values which resulted in 1,016 molecular descriptors for each compound.

### Feature Selection

We first used feature selection on the compounds in the proxy-DILI dataset using a variance threshold (as implemented in scikit-learn v1.1.1^66^) to filter features (Figure 1 Step 1). We used a low variance threshold of 0.05 for Morgan fingerprints resulting in 89 selected bits, a threshold of 0.10 for MACCS keys resulting in 100 selected keys, and a threshold of 0.10 for Mordred descriptors resulting in 653 selected descriptors. Lower thresholds for variance ensured strict selection criteria, leading to fewer selected features to strike a balance between the length of all fingerprints and physicochemical parameters. An additional 15 calculated physicochemical parameters (as implemented in RDKit^64^ v2022.09.5: topological polar surface area, hydrogen bond acceptors and donors, fraction of sp3 carbons, log P, and the number of rotatable bonds, rings, assembled rings, aromatic rings, hetero atoms, stereocenters, positive and negatively charged atoms, and the counts of NHOH and NO) were also added. This resulted in 189 bit-vector structural fingerprints and 668 molecular descriptors for each compound in the proxy-DILI dataset. The same selected features were used for the gold standard DILI dataset (Figure 1 Step 4) to avoid any information leaks.

### Evaluation of predictions from individual proxy-DILI models

First, we trained individual models for each of the nine proxy-DILI endpoints for all of the other proxy -DILI endpoints. For each proxy-DILI endpoint, we trained individual Random Forest models (Figure 1 Step 2) with a 5-fold stratified cross-validation and random halving search hyperparameter optimisation (as implemented in scikit-learn v1.1.1^66^ with hyperparameter space given in Supplementary Table S5). We used this hyperparameter-optimised model to obtain predicted probabilities for all compounds for the other proxy-DILI endpoints for every 9×9 combination. For each model built on a proxy-DILI endpoint, we chose an optimal decision threshold based on the J-statistic value (see released code for implementation) by comparing the predicted probabilities to the true values. We obtained final binary predictions using this threshold thereby choosing the best-case scenario where the balanced accuracy is optimised from the AUC-ROC curve. Next, we compared how well each proxy-DILI model was at predicting other proxy-DILI labels by comparing the F1 Score and Likelihood Ratios.

### Evaluating predictivity of individual proxy-DILI models for the gold standard DILI dataset

To train and evaluate models for DILI, we first split our gold-standard DILI dataset (containing 1,111 unique compounds) using ButinaSplitter based on the butina clustering of a bulk tanimoto fingerprint matrix split with a cutoff threshold of 0.70 (as implemented in DeepChem^70,71^, Figure 1 Step 3). This led to a training DILI data of 888 unique compounds (560 toxic and 328 non-toxic compounds) and a held-out DILI test set of 223 unique compounds (156 toxic and 67 non-toxic). This ensures that the DILI test set included a wide variety of compounds that are structurally less similar from those used in training. Therefore, the held-out DILI test set is a more challenging representation (compared to random splits) as encountered in real-world drug discovery: predicting DILI outcomes for new compounds away from the chemical space of the known compounds. We evaluated the performance of individual models built on each of the nine proxy-DILI endpoints on the held-out DILI test set (223 compounds). First, for each of the nine individual models, we obtained out-of-fold predicted probabilities on the DILI training data (888 compounds) using cross-validation with a 5-fold stratified split. We used these out-of-fold predicted probabilities and true values to obtain an optimal decision threshold based on the J-statistic value. Finally, we used each of the individual models and the corresponding optimal decision threshold to obtain predictions of the held-out DILI test set. We used the Jaccard similarity coefficient score (as implemented in scikit-learn v1.1.1^66^) to compare the similarity of predictions, that is, the predicted DILI vectors from each model. The Jaccard similarity coefficient measures the similarity between two sets of data counting mutual presence (positives/toxic) as matches but not the absences.

### Models for prediction of Cmax

Next, we trained two Random Forest regressor models to predict the median pMolar unbound plasma concentration and median pMolar total plasma concentration for 515 and 718 compounds respectively (Figure 1 Step 2). We used the selected 189 bit-vector structural fingerprints and 668 molecular descriptors as features to train the models with a 5-fold stratified cross-validation and random halving search hyperparameter optimisation as described above. The best estimator was refit on the entire dataset and the final model was used to generate predictions for compounds and these predicted features were used for training DILI models.

### Models for prediction of DILI

In this study, we built models (Figure 1 Step 5) using (a) selected 189-bit structural fingerprints, (b) selected 688 molecular descriptors, (c) a combination of selected 189-bit structural fingerprints and selected 668 molecular descriptors, (d) predicted nine proxy-DILI labels and two predicted pharmacokinetic parameters which refers to a FeatureNet approach, and (e) a combination of all three features spaces.

For each feature space, we used repeated nested cross-validation. First, the DILI training data was split into 5-folds. One of these folds was used as a validation set while the data from the remaining 4 folds were used to train and hyperparameter optimise a Random Forest Classifier (as implemented in scikit-learn v1.1.1^66^). We optimised the classifier model using a random halving search (as implemented in scikit-learn v1.1.1^66^) and 4-fold cross-validation (see Supplementary Table S5 for hyperparameter space). Once hyperparameters were optimised, we then used the fitted model to generate 4-fold cross-validated estimates for each compound in the fitted data. These predicted probabilities along with the real data were used to generate an optimal threshold using the J statistic value (see released code for implementation). Finally, we predicted the DILI endpoint for the validation set and used the optimal threshold to determine the DILI toxicity. The process was repeated 5 times in total until all 888 compounds in the DILI training data were used as a validation set. This entire nested-cross validation set-up was repeated ten times with different splits. The model with the highest AUC was fit on the entire DILI training data and we obtained the optimal threshold using the J statistic value on the 4-fold cross-validated estimates for each of these compounds. Finally, this threshold was used to evaluate our models (Figure 1 Step 6) on the held-out DILI test set (223 unique compounds). Thus, for each model using a feature space (or the combination), we obtained evaluation metrics on (a) the nested cross-validation (on training data), and (b) the held-out test set. The best-performing model (Figure 1 Step 7), as shown in the Results section, was the combination of all three feature spaces. This model was retrained (Figure 1 Step 8) on the complete gold-standard DILI dataset consisting of 1,111 distinct compounds. This model, DILIPredictor, can be accessed through a web application https://broad.io/DILIPredictor and have all code available for local use on GitHub at https://github.com/srijitseal/DILI.

To calculate the structural similarity of the held-out test to training data, we first calculated pairwise Tanimoto similarity (using 2048-bit Morgan fingerprint, see released code for implementation) for each test compound to each training compound. Finally, we calculated the mean of the three highest Tanimoto similarities (that is the three nearest neighbours) which was used to define the structural similarity of the particular test compound.

### Evaluation Metrics

All predictions (nested-cross validation and held-out test set) were evaluated using sensitivity, specificity, balanced accuracy (BA), Mathew’s correlation constant (MCC), F1 scores, positive predictive value (PPV), likelihood ratio (LR+)^72^, average precision score (AP), Area Under Curve-Receiver Operating Characteristic (AUC-ROC) as implemented in scikit-learn v1.1.1^66^.

### Feature importance measures to understand the chemistry and biological mechanisms for common DILI compounds

For the final model released publicly that used a combination of all feature spaces, we used SHAP values (as implemented in the shap python package^73^) to obtain feature importance for each input compound. This included proxy-DILI data, pharmacokinetic parameters, physicochemical features as well as MACCS key substructures that contributed to DILI toxicity/safety. Further, we show how DILIPredictor can be used to eluate the causes of DILI, both in chemistry and via mechanisms on the biological level using the importance measures on proxy-DILI labels. We analysed 4 compounds that were not present in the training data of these models. Two of these compounds, enzalutamide and sitaxentan, are known to cause DILI in humans while two compounds, 2-butoxyethanol and astaxanthin, did not cause DILI in humans. Additionally predicted profiles from another 12 compounds (also present in the training data) are shown in Supplementary Table S6. Several of toxic compounds were related to the study by Chang et al who compiled compounds causing DILI in patients undergoing chemotherapy.^74^ We also included two pairs of compounds studied by Chen et al such as doxycycline/minocycline and moxifloxacin/trovafloxacin; these pairs were defined by a similar chemical structure and mechanism of action but differed in their liver toxicity effects.^75^

### Statistics and Reproducibility

We have released the datasets used in this proof-of-concept study which are publicly available at https://broad.io/DILIPredictor. We released the Python code for the models which are publicly available on GitHub at https://github.com/srijitseal/DILI.

## RESULTS AND DISCUSSION

In this work, we trained models on each of nine proxy-DILI endpoints related to liver toxicity. We used these models to obtain predicted proxy-DILI labels for 1,111 compounds in the gold standard DILI dataset (as defined in Methods) none of which overlapped with the proxy-DILI dataset. We then trained new models using those predicted proxy-DILI labels as inputs, which refers to a FeatureNet approach, together with the compounds’ structural fingerprints, physicochemical properties, and a combination thereof, for 888 compounds the gold standard DILI datasets. We then evaluated the models on a held-out test set of 223 compounds.

### Comparing chemical spaces for the proxy-DILI and gold standard DILI datasets

We first examined the diversity and representation of compounds in the proxy-DILI and gold standard DILI datasets, to ensure the evaluation would be reasonable. The distribution of compounds in each of the nine labels of the proxy-DILI dataset covers a diverse range of physicochemical parameters as shown in Supplementary Figure S1. Gold standard DILI compounds effectively capture the diversity and representativeness of the compounds in the proxy-DILI dataset as shown in Supplementary Figure S2 for the physicochemical space of the 1,111 compounds in the gold standard DILI dataset compared to 12,133 compounds in the proxy-DILI dataset. Further, the held-out DILI test set (223 compounds) was also representative in the physicochemical parameter space of the training DILI data (888 compounds) as shown in Supplementary Figure S3. The main caveat to consider is that the six characteristic physicochemical descriptors capture the variability of physicochemical space only to a certain extent. Overall, we conclude that the chemical space covered by the datasets is sufficiently similar for our evaluation to be reliable.

### Concordance of Proxy-DILI dataset and DILI compounds

Next, we aimed to evaluate the concordance of labels in the proxy-DILI dataset with the gold standard DILI dataset. To do so, we compared all 13,703 compounds in the proxy-DILI dataset to the 1,111 compounds in the gold standard DILI dataset. It is important to note that these compounds (that overlapped between the proxy-DILI and gold standard DILI dataset) were only used to analyse concordance in this section and not in training the models, because that would leak information. As depicted in Figure 2, we observed a strong concordance between the data sourced from human hepatotoxicity dataset and preclinical data (Cohen’s Kappa = 0.60), and the three diverse DILI datasets (0.49 and 0.55) used in this study. The lack of perfect concordance is reasonable given these datasets are primarily derived from human-related data, as opposed to animal data or *in vitro* assays. Note, concordance between DILI and proxy-DILI labels may be affected as the proxy-DILI dataset used here includes some of the DILI compounds (these overlapped compounds were removed later when training models).

**Figure 2:**
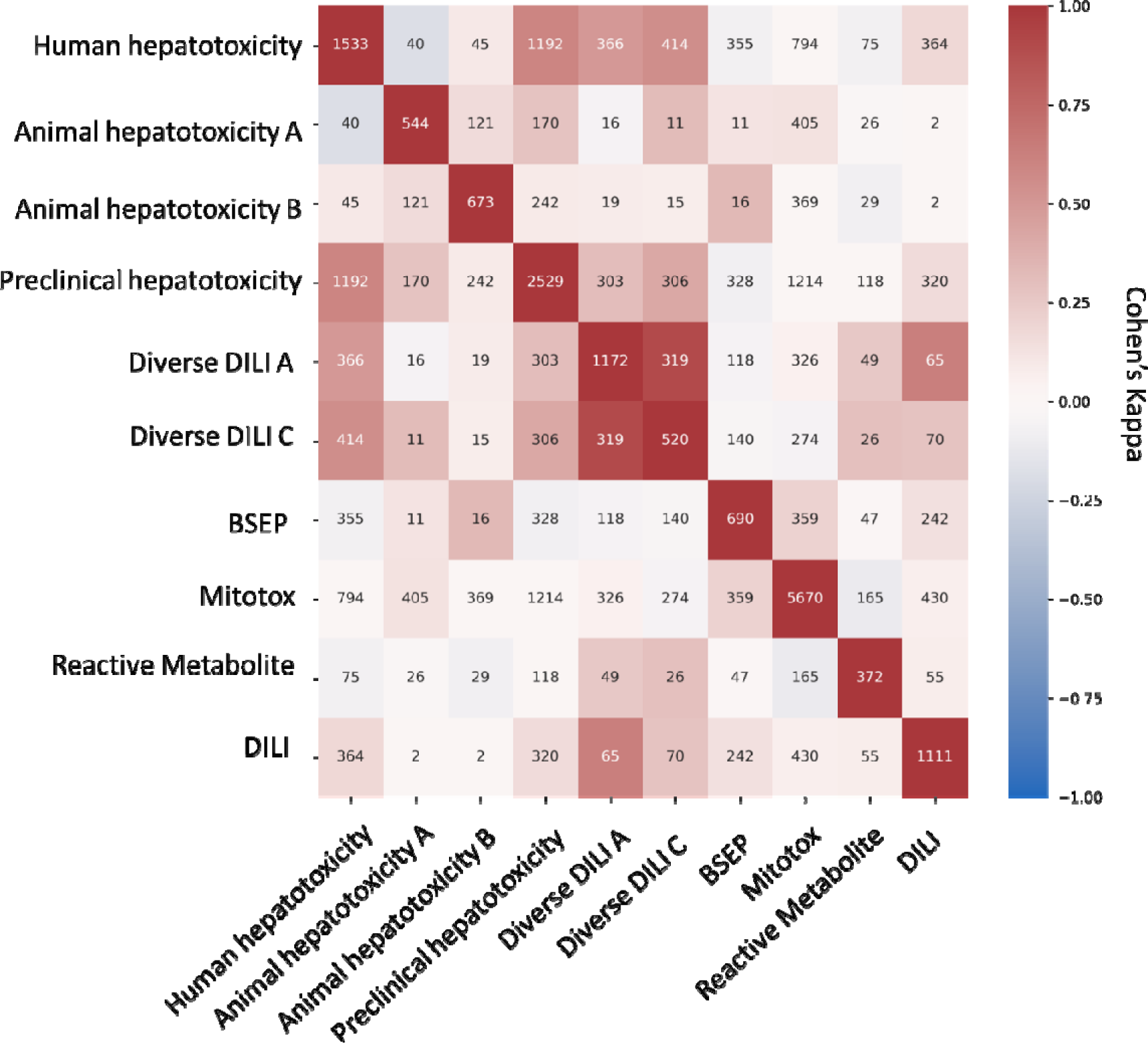
Concordance of compounds overlapping in-between nine labels in the proxy-DILI dataset (13,703 compounds) including compounds that overlapped with DILI data (1,111 compounds). Concordance is given using Cohen’s kappa (and the number of overlapping compounds given as annotations). Overall, the human-related proxy-DILI labels and diverse heterogenous DILI labels showed high concordance with DILI compounds and among each other.

### Individual proxy-DILI models are complementary to each other and distinct in their prediction for DILI compounds

We next used the individual models built on the nine proxy-DILI labels to predict the other proxy-DILI labels (with evaluation metrics as shown in Supplementary Table S7). As shown in Figure 3, we observed the human hepatotoxicity was well predicted using preclinical hepatotoxicity (LR+ = 3.63, F1=0.79). Bile salt export pump inhibition (BESP) and mitochondrial toxicity were strongly predictive of each other (LR+ = 2.16, F1= 0.36 when using BSEP to predict mitotox and LR+ = 3.35, F1=0.77 when using mitotox to predict BSEP). Overall, the assays in the proxy-DILI dataset can be used to train individual models to generate predicted proxy-DILI labels which then provide a complementary source of information.

**Figure 3.**
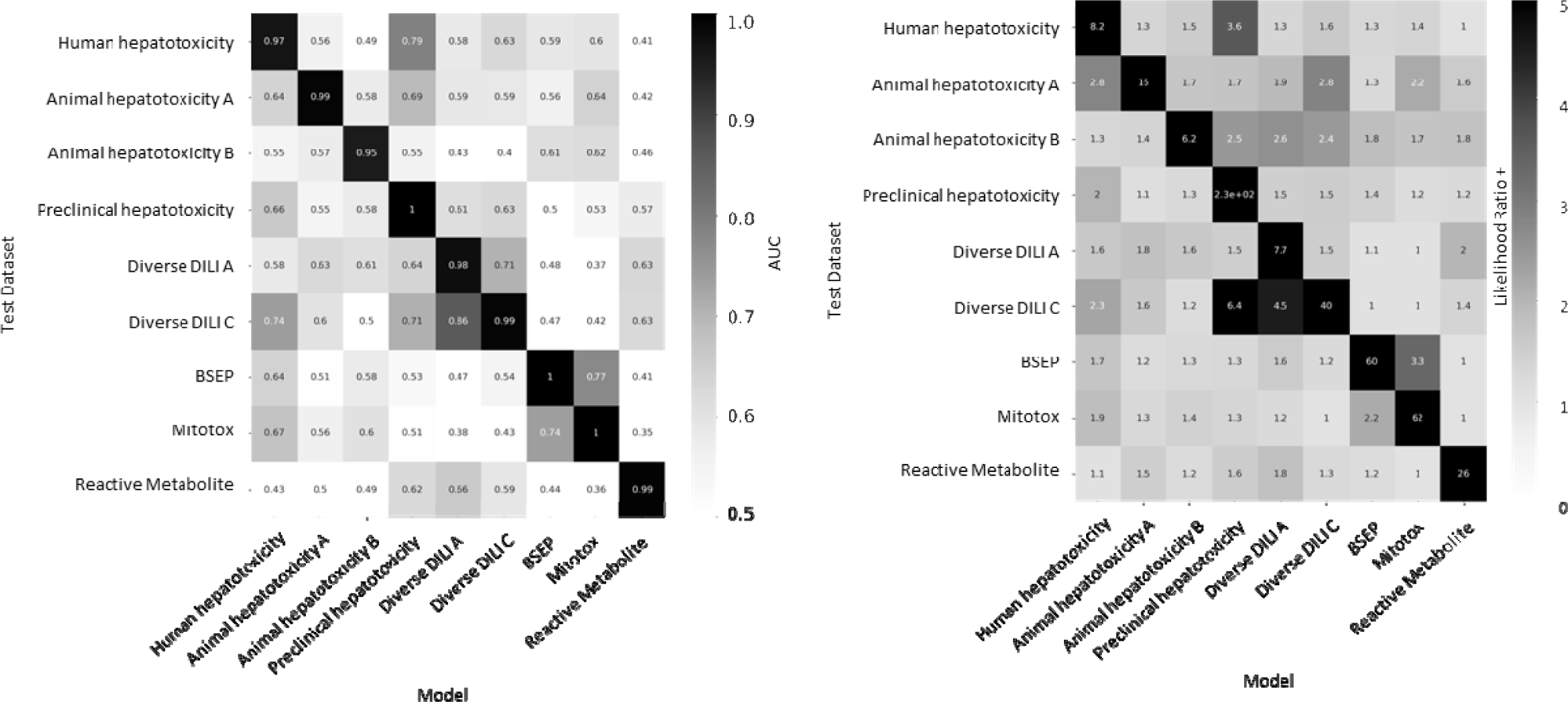
Performance metrics for models built on nine proxy-DILI labels when predicting labels for the other proxy-DILI in the model were evaluated using (a) AUC-ROC and (b) Likelihood Ratio (LR+).

We next analysed the nine individual proxy-DILI models and a model built on the two PK parameters (Cmax unbound and total) for their predictions on the 223 compounds in the held-out compounds of the gold standard DILI dataset. As shown in Figure 4 (further details in Supplementary Table S8), the best-performing models were the model built on the preclinical animal hepatotoxicity (AUC=0.61, LR+ = 1.63) and the model built on diverse DILI C dataset (AUC=0.59, LR+ = 1.32). Biological labels have compounds covering a wider biological and chemical space coverage which warrants their inclusion in our study as shown by Jaccard similarity for predictions on the held-out DILI dataset. Predictions from models built on animal hepatotoxicity labels were not similar to predictions from models built on human hepatotoxicity labels (Figure 5; mean Jaccard similarity of 0.12). We found that predictions from models built on human-related labels were similar (e.g., predictions from the preclinical hepatotoxicity model have a Jaccard similarity of 0.42). However, predictions from human-related labels were dissimilar to predictions from *in vitro* assays (e.g., predictions from the preclinical hepatotoxicity model had only a 0.04 Jaccard similarity to predictions from the mitotox model and 0.02 Jaccard similarity to predictions from the reactive metabolite formation model). Overall, we conclude that each model built on a proxy-DILI label and the PK parameters was distinctive in its prediction, thus providing complementary information on compounds’ potential for DILI.

**Figure 4.**
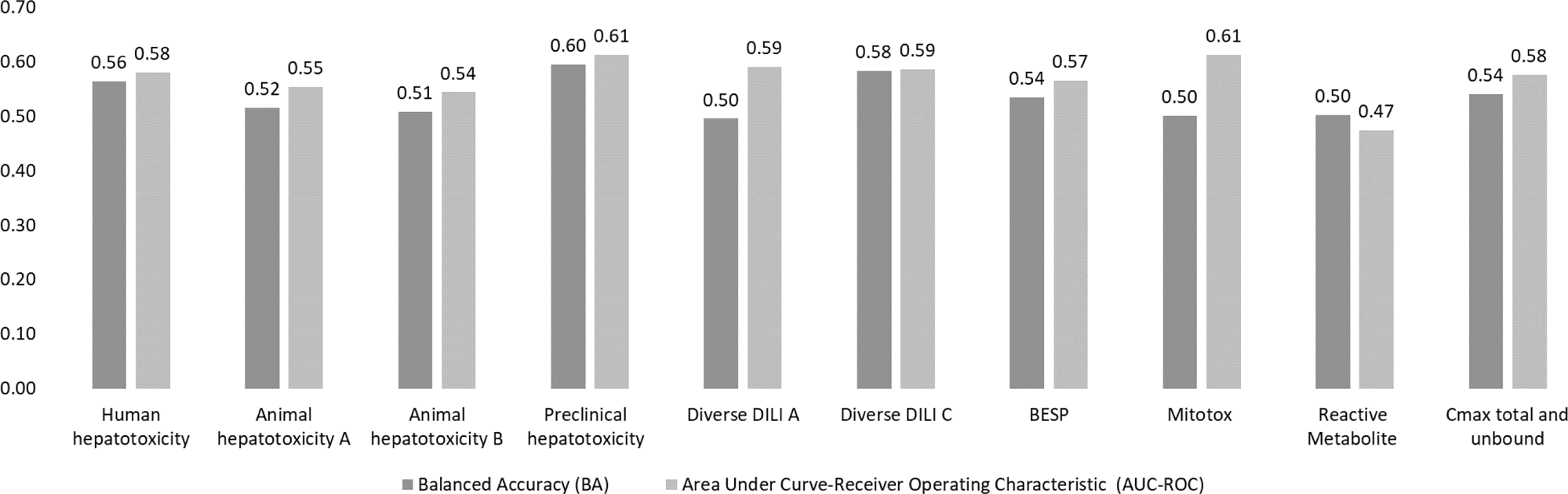
Performance Metrics AUC-ROC and Balanced Accuracy achieved by each of nine individual models built on the proxy-DILI labels and a model built on two pharmacokinetic parameters (Cmax total and unbound) when tested on the 255 compounds in the held-out DILI dataset.

**Figure 5:**
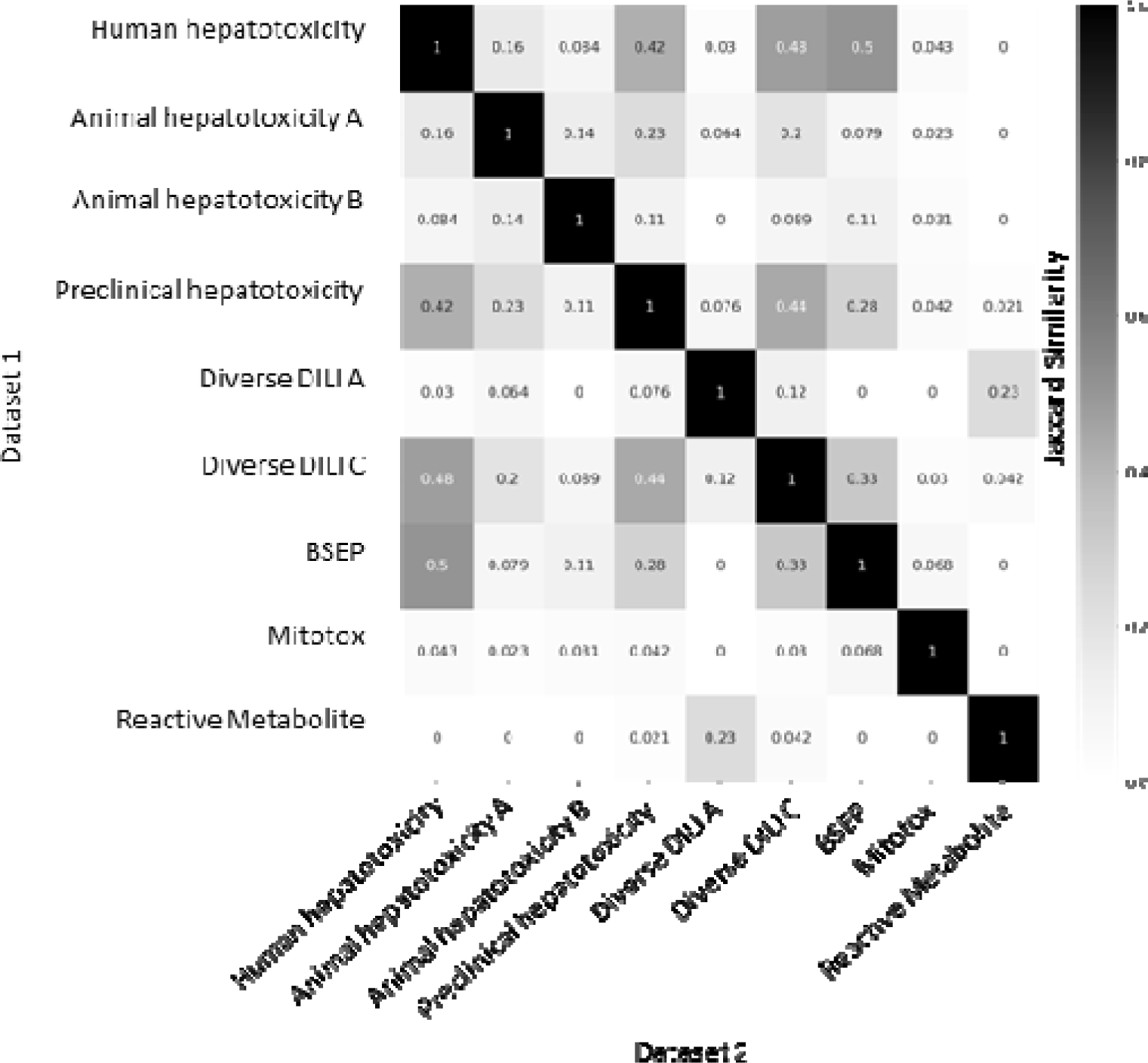
Jaccard Similarity of predictions on the held-out DILI dataset (223 compounds) for individual models built on nine proxy-DILI labels in the proxy-DILI dataset.

### Models combining chemical structure, physicochemical properties, PK parameters and predicted proxy-DILI data outperform individual models

We next compared models built on combinations of proxy-DILI labels (including PK parameters), chemical structure, and physicochemical properties including Mordred descriptors (Table 2). When comparing results from 55 held-out test sets from the repeated nested cross-validation (as shown in Figure 6 with the comparison of differences in distribution using a paired t-test.), the models combining structural fingerprints, physicochemical properties, Mordred descriptors, PK parameters and predicted proxy-DILI labels achieved a mean balanced accuracy (BA) of 0.64 (mean LR+ = 1.84) compared to the second-best models using only physicochemical properties and Mordred descriptors with a mean BA of 0.64 (mean LR+ = 1.83) and models using structural fingerprints, physicochemical properties and Mordred descriptors which also achieved a mean BA of 0.63 (mean LR+ = 1.81). Models using only structural fingerprints achieved a mean BA of 0.63 (mean LR+ = 1.78) while models using only predicted proxy-DILI labels and PK parameters as features achieved a mean BA of 0.61 (mean LR+ = 1.77) in the nested cross-validation. Supplementary Figure S4 compares the distribution of positive predictive value for all model combinations using all feature sets (predicted proxy-DILI labels and PK parameters, structural features, and Mordred physicochemical descriptors).

**Figure 6.**
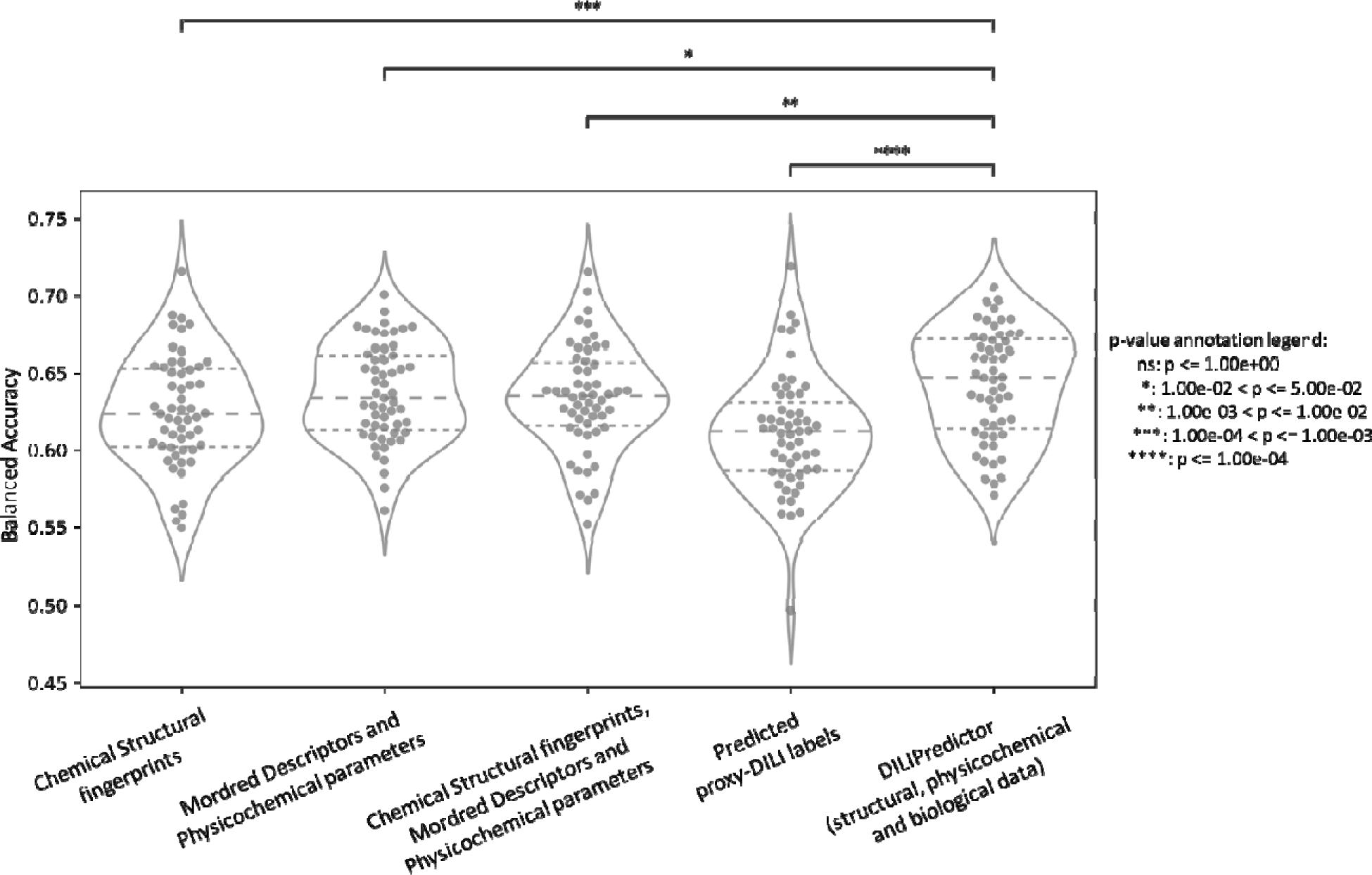
Performance metrics Balanced Accuracy for combination models from 55 held-out test sets from repeated nested cross-validation using (a) selected 189-bit structural fingerprints, (b) selected 668 molecular descriptors, (c) selected 189-bit structural fingerprints and selected 668 molecular descriptors, (d) predicted nine proxy-DILI labels and 2 PK parameters, and (e) a combination of all three features spaces, compared with a paired t-test.

**Table 2:**
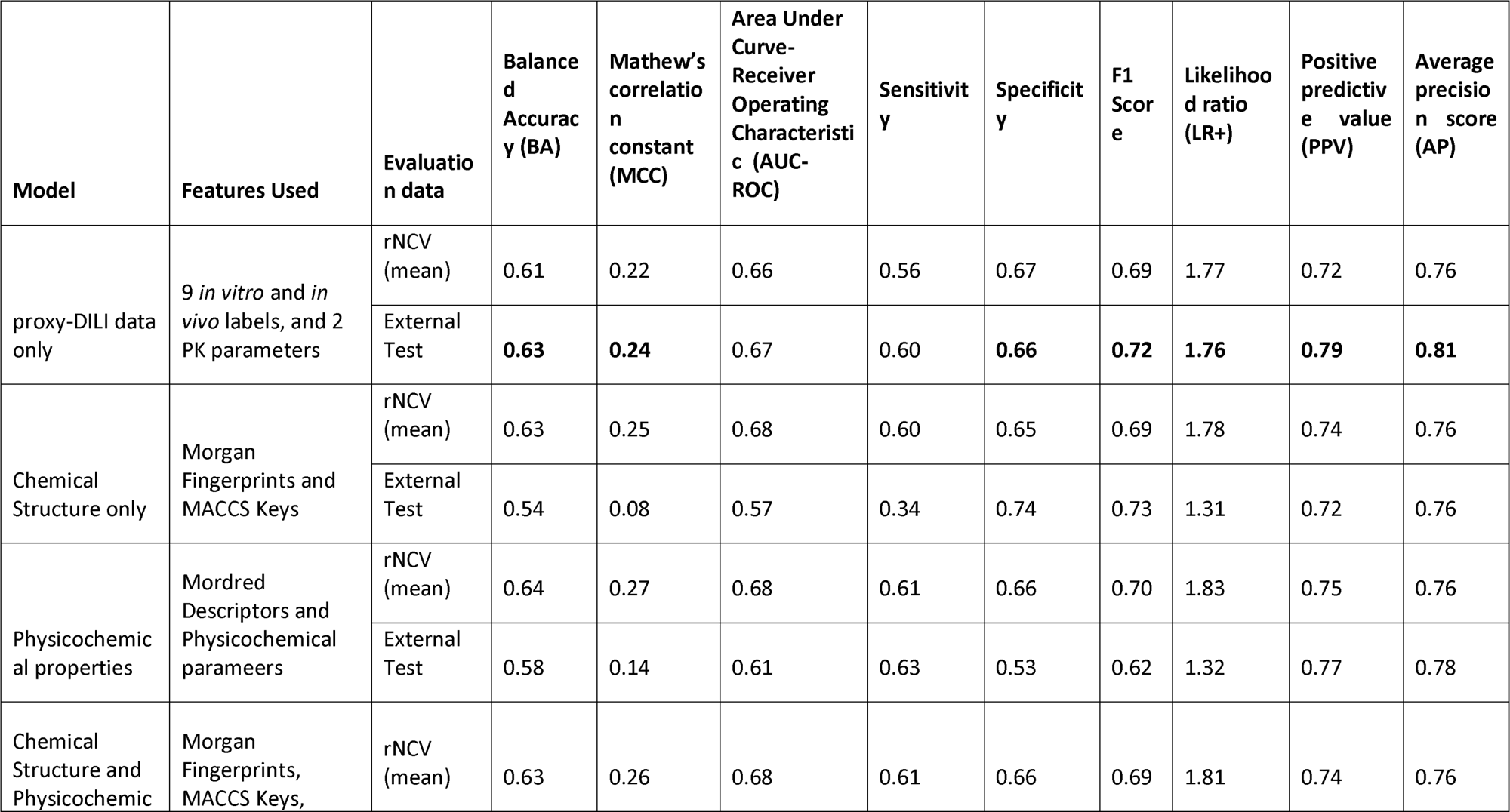
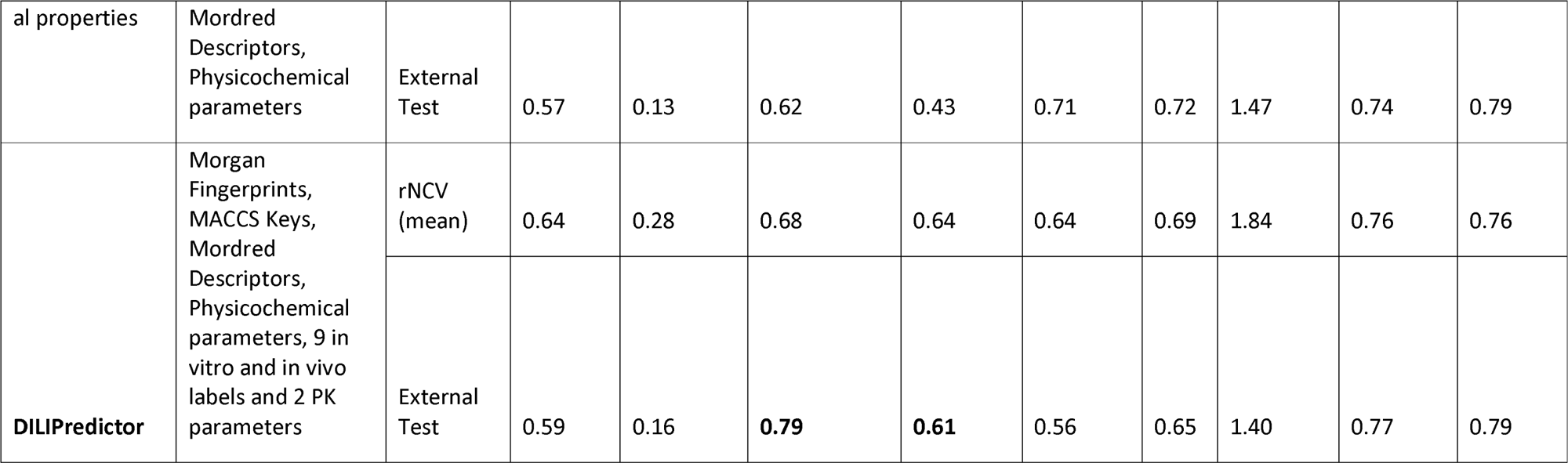
Performance of various models from (a) 55 held out test sets from repeated nested cross validation and (b) for the 223 compounds in the held-out DILI dataset. rNCV: repeated nested cross validation.

We next retrained all hyperparameter-optimised models on the DILI training data (888 compounds) and evaluated the final models on the held-out DILI test set (223 compounds). The DILIPredictor model (combining all predicted proxy-DILI labels and PK parameters, structural features, and Mordred physicochemical descriptors) achieved an AUC = 0.79 (LR+ = 1.40) (Table 2). The model using only proxy-DILI and PK parameters achieved an AUC = 0.67 (LR+ = 1.76). Other models achieved AUC = 0.62 (LR+ = 1.47) using structural, Mordred and physicochemical descriptors, AUC = 0.54 (LR+ = 1.31) using chemical structural only, and AUC = 0.61 (LR+ = 1.32) for the model using Mordred and physicochemical descriptors.

One metric relevant in predictive safety/toxicology is the positive likelihood ratio^72^ and the detection of toxic compounds with a lower false positive rate. Improved detection with lower false positive rates aids in evaluating model performance across various threshold settings, shifting the focus from AUC as a singular statistical value to a more nuanced examination along the AUC-ROC curve from a false positive rate of 0 to 1. When predicting the first 29 compounds correctly as true positives (or approximately 13% of the 223 compounds in the held-out test set), DILIPredictor achieved the highest LR+ score of 2.68 (25 toxic compounds correctly predicted out of 29 compounds, PPV = 0.86) compared to the structural model which achieved an LR+ score of 1.65 (23 toxic compounds correctly predicted out of 29 compounds, PPV = 0.78). This improvement is mainly from being able to detect compounds at a wider range of structural similarity to training data (as shown in Supplementary Figure S5 using the distribution of the top true positives detected with low false positive rates for each model). Overall, this shows that using all feature types in DILIPredictor allows for the detection of a greater number of toxic compounds with a low false positive rate.

We subsequently compared our models to those reported in earlier publications. Table 3 presents a selection of recent DILI prediction models that employ chemical features and biological data to predict liver toxicity. Since most previous studies did not emphasize likelihood ratios, and often not scaffold-based splits, it is not possible to compare LR+ scores; therefore, we can only make comparisons within the models developed in this study. It is important to note that the size, source, and consequently the quality of training and test datasets vary across previous literature, rendering direct comparisons infeasible. In our study, the final DILIPredictor model achieved a high AUC-ROC of 0.79 on the held-out gold standard DILI dataset (223 compounds), which aligns with the average AUC (0.73) from prior studies. The balanced accuracy of DILIPredictor in this study, standing at 0.59, is lower compared to previous models, averaging 0.64. This difference is likely caused due to a stringent scaffold-based split for the external test set, which decreases the numerical score but better mirrors the real-world drug discovery process. We also adopted a fixed held-out test set that is scaffold-split, in contrast to less rigorous random splits and external datasets used in other studies (which contain some similar compounds to training data). Furthermore, we delved beyond the statistical value of AUC-ROC to examine likelihood ratios and improved detection with low false positive rates. This approach allows us to evaluate the quality of the AUC-ROC curve rather than reducing it to a single statistical value.

**Table 3:** Previously published models used in the evaluation of hepatotoxicity/liver injury (for test sets only)

| Model | Features | Compounds in Train set | Compounds in Test set | Test Splitting Strategy | Balanced Accuracy | AUC-ROC |
| --- | --- | --- | --- | --- | --- | --- |
| Ensemble of RF and SVM | Molecular fingerprints | 1241 | 286 | External Dataset | 0.82 | 0.9 |
| Random Forests | Imaging Phenotypes and Chemical Descriptors | 346 | 41 | Literature Survey | 0.52 | 0.74 |
| Ensemble Models | Molecular features, physicochemical properties | 1254 | 204 | Literature Survey | 0.72 | 0.73 |
| Random Forests | 2D molecular descriptors | 996 | 341 | External Dataset | 0.67 | 0.71 |
| Random Forests | 2D molecular descriptors | 996 | 921 | External Dataset | 0.57 | 0.59 |
| SVM | Morgan Fingerprints | 923 | 49 | External Dataset | 0.67 | - |
| SVM | Predicted protein targets | 923 | 49 | External Dataset | 0.59 | - |
| Random Forests | 0–2.5D molecular descriptors | 1075 | 554 | External Dataset | 0.77 | 0.81 |
| GA-SVM | 2D and 3D molecular descriptors | 3712 | 269 | Proprietary Dataset | 0.64 | 0.68 |
| Random Forests | Morgan Fingerprints | 845 | 362 | Random Splits (repeated 100 times) | - | 0.75 |
| Naïve Bayes | Morgan Fingerprints | 336 | 84 | Random Split | 0.73 | 0.81 |
| SVM | MACCS keys | 1317 | 88 | External Dataset | 0.68 | 0.62 |
| Rule-Based | Physicochemical properties and common toxicity mechanisms | - | 200 | N/A | 0.32 | - |
| **Average** |  |  |  |  | **0.64** | **0.73** |
| Random Forests | Structural, Physicochemical, predicted in vitro, in vivo and PK parameters | 888 | 223 | Scaffold-split | **0.59** (scaffold-split) | **0.79** (scaffold-split) |

### Feature Interpretation

We next used feature interpretation to analyse the chemistry and biological mechanisms for compounds known to cause DILI. We chose four compounds (Table 4) of which two were known for their DILI^74^ (namely, enzalutamide and sitaxentan) and two compounds that do not cause DILI in humans (namely, 2-butoxyethanol and astaxanthin). DILIPredictor could detect structural information relevant to causing DILI (four compounds shown in Figure 7 and further in Table 4 and Supplementary Figure S6). As shown in Figure 7, sitaxentan (a sulfonamide-based ETA receptor antagonist) was predicted toxic, with a positive contribution from the MACCS substructure near the sulphonamide, which is known to cause human liver injuries.^76^ The MACCS features most contributing to the toxicity for paclitaxel, docetaxel, and cabazitaxel, (which contain a taxane group as shown in Supplementary Figure S6) was found to be the presence of a taxadiene core. These compounds stabilise microtubules by binding to the β-tubulin and are known to cause mitochondrial toxicity.^77^ Further, DILIPredictor correctly predicted compounds such as 2-butoxyethanol and astaxanthin to be non-toxic in humans even though they cause hepatic injury in animal models^22,30^ (Figure 7). In 2-butoxyethanol, proxy-DILI features associated with either animal hepatotoxicity or preclinical hepatotoxicity contributed to predicting toxicity in humans, however, the proxy-DILI indicators related to human hepatotoxicity ultimately led to the prediction of non-toxicity.

**Figure 7:**
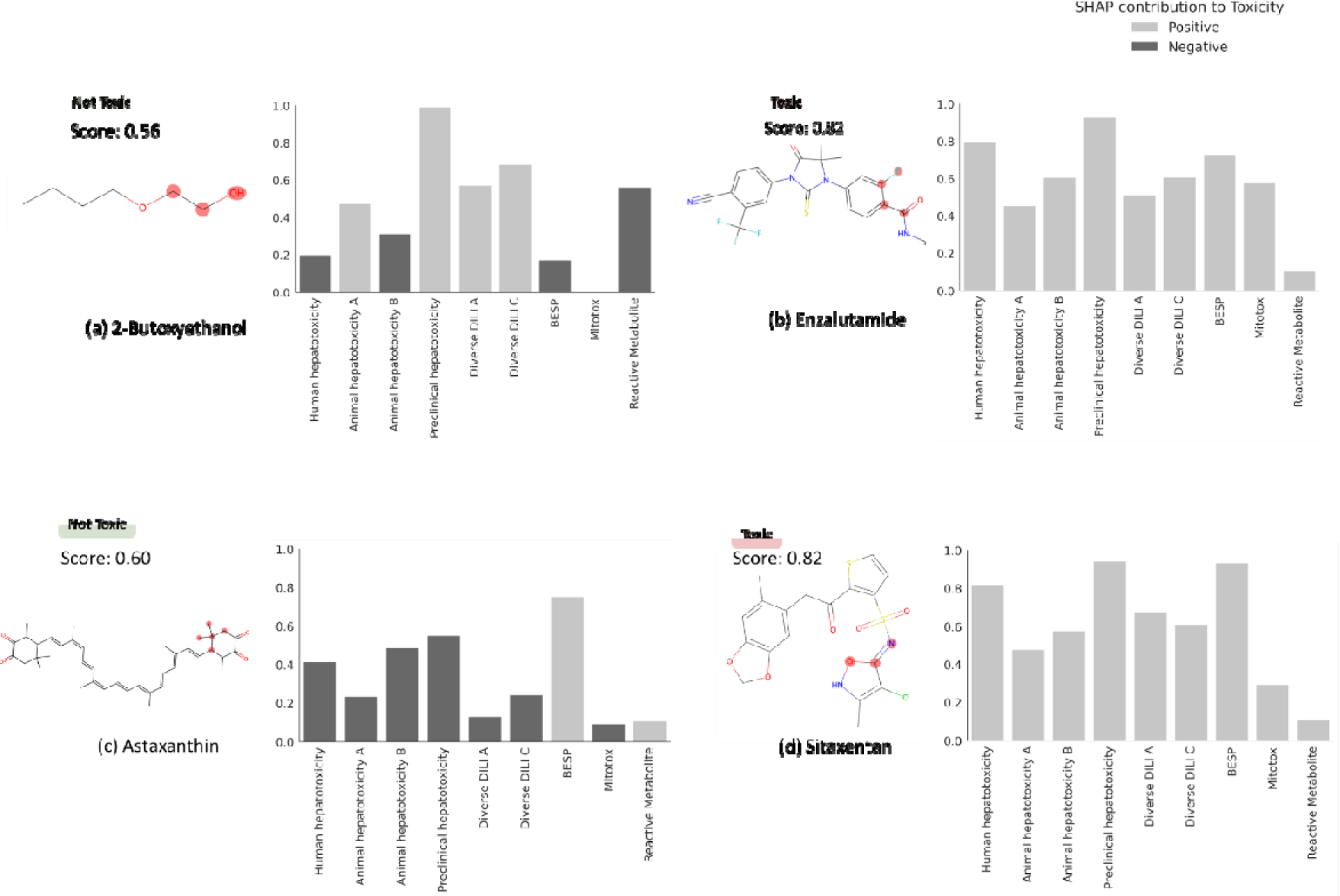
MACCS substructure (highlighted) and proxy-DILI labels contributing to DILI when using DILIPredictor (SHAP values) for two compounds known to cause DILI and for two compounds which do not cause DILI in humans (further details in Table 4, and for another 12 compounds in Supplementary Figure S6 and Supplementary Table S6). The highest contribution to toxicity/safety from the MACCS substructure is highlighted with the chemical structure.

**Table 4:** DILI predictions for 14 compounds known to cause DILI and 2 compounds which do not cause DILI in humans (not used in training models in this study) and top 3 proxy-DILI labels positively and negatively for contributing to the prediction.

| Compound Name | DILI (Literature) | DILI Prediction | DILI Probability | Remarks | Most Contribution proxy-DILI endpoints to Prediction |  |  |
| --- | --- | --- | --- | --- | --- | --- | --- |
|  |  |  |  |  | Ranked 1 | Ranked 2 | Ranked 3 |
| 2-Butoxyethanol | Not Toxic | Not Toxic | 0.56 | Known DILI in mice, not in human | Mitotox | Human hepatotoxicity | Animal hepatotoxicity B |
| Astaxanthin | Not Toxic | Not Toxic | 0.6 | Vitamin A derivative; High structural similarity to retinoid but does not cause DILI | Diverse DILI A | Preclinical hepatotoxicity | Diverse DILI C |
| Enzalutamide | Toxic | Toxic | 0.82 |  | Preclinical hepatotoxicity | Human hepatotoxicity | Mitotox |
| Sitaxentan | Toxic | Toxic | 0.82 | Withdrawn | Preclinical hepatotoxicity | Human hepatotoxicity | Mitotox |

Finally, among structurally similar pairs of compounds, acitretin was correctly predicted as toxic while astaxanthin was correctly predicted to be non-toxic. For acitretin, the preclinical hepatotoxicity label contributed to the toxicity prediction. Conversely, labels associated with human hepatotoxicity contributed to correctly predicting astaxanthin as non-toxic. Among tetracyclines, pairs of compounds doxycycline (prediction scores = 0.66) and minocycline (0.66), and among fluoroquinolones, pairs of compounds moxifloxacin (0.74) and trovafloxacin (0.84) were correctly predicted toxic. For fluoroquinolones, the prediction scores obtained from DILIPredictor were in agreement with the less-toxic or more-toxic DILI annotations collated by Chen et al.^75^ Among compounds withdrawn from market, sitaxentan and trovafloxacin were flagged with prediction scores above 0.80 threshold; many compounds currently on the market such as docetaxel and paclitaxel were also flagged in the 0.70 to 0.75 threshold as being DILI-toxic. Overall, DILIPredictor combined chemical structures and biological data to correctly predict DILI in humans.

### Limitations and Future Directions

The primary focus of this study was the generation of binary classification models for drug-induced liver injury, and this is empirically based on known datasets. Like most previously published models, empirical models can still be very useful for decision making, provided they are well predictive, in particular for novel chemical space.^78,79^ Besides using predicted Cmax (unbound ant total), we did not incorporate factors such as dose or time point into this study due to its scarcity in available public data. Labelling schemes are not always binary but sometimes include an “ambiguous” class (such as in the DILIrank dataset) and these compounds are hence not included in this study. While *in vitro* data can provide valuable insights into drug toxicities, they are still proxy endpoints for the *in vivo* effects. Toxic compounds detected in *in vitro* assays can often cause corresponding toxicity *in vivo*, but compounds that appear to be safe in *in vitro* are not necessarily safe in humans.^28,29^ In the future, with the availability of larger relevant -omics datasets, such as from the Omics for Assessing Signatures for Integrated Safety Consortium (OASIS) Consortium^80^, multitask learning models such as Conditional Neural Processes can be implemented which can train on several predictive tasks simultaneously by shared representation learning.^81^ In the future, with the availability of larger -omics datasets and histopathology readouts, it could be possible to explore mechanistic underpinnings regarding Molecular Initiating Events or Key Events pathways of toxicity.

## CONCLUSIONS

In this work, we trained models to predict drug-induced liver injury (DILI) using not only chemical data but also heterogeneous biological *in vivo* (human and animal) and *in vitro* data from various sources. We found a strong concordance in observed data between compounds with the proxy-DILI labels and DILI compounds. The nine proxy-DILI models were not predictive of each other-this complementarity suggests that they could be combined to predict drug-induced liver injury. Random Forest models that combined different types of input data - structural fingerprints, physicochemical properties, PK properties and proxy-DILI labels - improved predictive performance, especially in detection with low false positive rates, with the highest LR+ score of 2.68 (25 toxic compounds with PPV=0.93). DILIPredictor accurately predicted the toxicity of various compounds known to cause DILI, including fourteen notorious DILI-inducing compounds, by recognizing chemical structure as well as biological mechanisms. DILIPredictor was further able to differentiate between animal and human sensitivity for DILI and exhibited a potential for mechanism evaluation for these compounds. DILIPredictor was trained on existing in vitro and in vivo data. Using only chemical structures as the input and a FeatureNet approach, it can predict DILI risk more accurately, thus aiding decision-making for new compounds before new in vivo toxicology and pharmacokinetic data is collected (which typically involves animal experiments that we aim to reduce).DILIPredictor can also be integrated into Design-Make-Test-Analyse (DMTA) cycles to aid in the selection and modification of compounds before more extensive and expensive testing is conducted.^82^ Overall, the study demonstrated that incorporating all complementary sources of information can significantly improve the accuracy of DILI prediction models. Furthermore, the availability of larger, high-quality, and standardized datasets for DILI in the public domain can greatly enhance the development of predictive models for drug-induced liver injury such as from the Omics for Assessing Signatures for Integrated Safety Consortium (OASIS).^80^ We released our final interpretable models at (with all code available for download at GitHub at https://github.com/srijitseal/DILI) and datasets used in this study at https://broad.io/DILIPredictor. Further, DILI Predictor is available for direct implementations via https://pypi.org/project/dilipred/ and installable via ‘pip install dilipred’.

## ACKNOWLEDGEMENTS

S.S. acknowledges funding from the Cambridge Commonwealth, European and International Trust, Boak Student Support Fund (Clare Hall), Jawaharlal Nehru Memorial Fund, Allen, Meek and Read Fund, and Trinity Henry Barlow (Trinity College). O.S. acknowledges funding from the Swedish Research Council (grants 2020-03731 and 2020-01865), FORMAS (grant 2022-00940), Swedish Cancer Foundation (22 2412 Pj), and Horizon Europe grant agreement #101057014 (PARC) and #101057442 (REMEDI4ALL). S.S. and A.E.C. acknowledges funding from the National Institutes of Health (NIH MIRA R35 GM122547 to A.E.C.), the Massachusetts Life Sciences Center Bits to Bytes Capital Call program for funding the data production (to Shantanu Singh, Broad Institute of MIT and Harvard), and the OASIS Consortium organised by HESI. This work was performed using resources provided by the Cambridge Service for Data-Driven Discovery (CSD3) operated by the University of Cambridge Research Computing Service (www.csd3.cam.ac.uk), provided by Dell EMC and Intel using Tier-2 funding from the Engineering and Physical Sciences Research Council (capital grant EP/P020259/1), and DiRAC funding from the Science and Technology Facilities Council (www.dirac.ac.uk).

## ASSOCIATED CONTENT

**Supplemental Information**. The Supporting Information is available. We released the Python code for our models which are publicly available at https://github.com/srijitseal/DILI. The datasets are also uploaded on https://doi.org/10.5281/zenodo.11522509. The online tools are accessible at https://broad.io/DILIPredictor. DILI Predictor is also installable with ‘pip install dilipred’ (https://pypi.org/project/dilipred/).

## AUTHOR INFORMATION

All authors have approved the final version of the manuscript.

## Contributions

SS designed and performed data analysis, implemented, and trained the models. SS analysed the interpretation of features and the results of models. MM assisted with packaging and implementing dilipred for local implementation. SS wrote the manuscript with extensive discussions with OS and AB and who supervised the project. All the authors (SS, DPW, LHG, MM, AEC, OS, and AB) reviewed, edited, contributed to discussions, and approved the final version of the manuscript.

## REFERENCES

(1) Andrade, R. J.; Chalasani, N.; Björnsson, E. S.; Suzuki, A.; Kullak-Ublick, G. A.; Watkins, P. B.; Devarbhavi, H.; Merz, M.; Lucena, M. I.; Kaplowitz, N.; Aithal, G. P. Drug-Induced Liver Injury. Nat. Rev. Dis. Prim. 2019, 5 (1), 1–22. 10.1038/s41572-019-0105-0.

(2) Remmer, H. The Role of the Liver in Drug Metabolism. Am. J. Med. 1970, 49 (5), 617–629. 10.1016/S0002-9343(70)80129-2.

(3) Licata, A. Adverse Drug Reactions and Organ Damage: The Liver. Eur. J. Intern. Med. 2016, 28, 9–16. 10.1016/j.ejim.2015.12.017.

(4) Ostapowicz, G.; Fontana, R. J.; Schioødt, F. V.; Larson, A.; Davern, T. J.; Han, S. H. B.; McCashland, T. M.; Shakil, A. O.; Hay, J. E.; Hynan, L.; Crippin, J. S.; Blei, A. T.; Samuel, G.; Reisch, J.; Lee, W. M.; Santyanarayana, R.; Caldwell, C.; Shick, L.; Bass, N.; Rouillard, S.; Atillasoy, E.; Flamm, S.; Benner, K. G.; Rosen, H. R.; Martin, P.; Stribling, R.; Schiff, E. R.; Torres, M. B.; Navarro, V.; McGuire, B.; Chung, R.; Abraczinskas, D.; Dienstag, J. Results of a Prospective Study of Acute Liver Failure at 17 Tertiary Care Centers in the United States. Ann. Intern. Med. 2002, 137 (12), 947–954. 10.7326/0003-4819-137-12-200212170-00007.

(5) Onakpoya, I. J.; Heneghan, C. J.; Aronson, J. K. Post-Marketing Withdrawal of 462 Medicinal Products Because of Adverse Drug Reactions: A Systematic Review of the World Literature. BMC Med. 2016, 14 (1), 1–11. 10.1186/s12916-016-0553-2.

(6) Raschi, E.; De Ponti, F. Strategies for Early Prediction and Timely Recognition of Drug-Induced Liver Injury: The Case of Cyclin-Dependent Kinase 4/6 Inhibitors. Front. Pharmacol. 2019, 10 (OCT), 1235. 10.3389/fphar.2019.01235.

(7) Weaver, R. J.; Blomme, E. A.; Chadwick, A. E.; Copple, I. M.; Gerets, H. H. J.; Goldring, C. E.; Guillouzo, A.; Hewitt, P. G.; Ingelman-Sundberg, M.; Jensen, K. G.; Juhila, S.; Klingmüller, U.; Labbe, G.; Liguori, M. J.; Lovatt, C. A.; Morgan, P.; Naisbitt, D. J.; Pieters, R. H. H.; Snoeys, J.; van de Water, B.; Williams, D. P.; Park, B. K. Managing the Challenge of Drug-Induced Liver Injury: A Roadmap for the Development and Deployment of Preclinical Predictive Models. Nat. Rev. Drug Discov. 2020, 19 (2), 131–148. 10.1038/s41573-019-0048-x.

(8) Mihajlovic, M.; Vinken, M. Mitochondria as the Target of Hepatotoxicity and Drug-Induced Liver Injury: Molecular Mechanisms and Detection Methods. Int. J. Mol. Sci. 2022, 23 (6). 10.3390/ijms23063315.

(9) Kenna, J. G.; Taskar, K. S.; Battista, C.; Bourdet, D. L.; Brouwer, K. L. R.; Brouwer, K. R.; Dai, D.; Funk, C.; Hafey, M. J.; Lai, Y.; Maher, J.; Pak, Y. A.; Pedersen, J. M.; Polli, J. W.; Rodrigues, A. D.; Watkins, P. B.; Yang, K.; Yucha, R. W. Can Bile Salt Export Pump Inhibition Testing in Drug Discovery and Development Reduce Liver Injury Risk? An International Transporter Consortium Perspective. Clin. Pharmacol. Ther. 2018, 104 (5), 916–932. 10.1002/cpt.1222.

(10) Villanueva-Paz, M.; Morán, L.; López-Alcántara, N.; Freixo, C.; Andrade, R. J.; Lucena, M. I.; Cubero, F. J. Oxidative Stress in Drug-Induced Liver Injury (DILI): From Mechanisms to Biomarkers for Use in Clinical Practice. Antioxidants 2021, 10 (3), 1–35. 10.3390/antiox10030390.

(11) Bao, Y.; Wang, P.; Shao, X.; Zhu, J.; Xiao, J.; Shi, J.; Zhang, L.; Zhu, H. J.; Ma, X.; Manautou, J. E.; Zhong, X. B. Acetaminophen-Induced Liver Injury Alters Expression and Activities of Cytochrome P450 Enzymes in an Age-Dependent Manner in Mouse Liver. Drug Metab. Dispos. 2020, 48 (5), 326–336. 10.1124/DMD.119.089557.

(12) Johansson, J.; Larsson, M. H.; Hornberg, J. J. Predictive in Vitro Toxicology Screening to Guide Chemical Design in Drug Discovery. Curr. Opin. Toxicol. 2019, 15, 99–108. 10.1016/j.cotox.2019.08.005.

(13) Biomea Fusion Announces BMF-219 in Diabetes Placed on https://www.globenewswire.com/news-release/2024/06/06/2895065/0/en/Biomea-Fusion-Announces-BMF-219-in-Diabetes-Placed-on-Clinical-Hold.html (accessed Jun 7, 2024).

(14) RAPT Therapeutics Announces Clinical Hold on Studies https://www.globenewswire.com/news-release/2024/02/20/2831721/0/en/RAPT-Therapeutics-Announces-Clinical-Hold-on-Studies-Evaluating-Zelnecirnon.html (accessed Jun 7, 2024).

(15) Tango Therapeutics Announces Discontinuation of TNG348 Program | Business Wire https://www.businesswire.com/news/home/20240523266195/en/Tango-Therapeutics-Announces-Discontinuation-of-TNG348-Program (accessed Jun 7, 2024).

(16) Rathman, J.; Yang, C.; Ribeiro, J. V.; Mostrag, A.; Thakkar, S.; Tong, W.; Hobocienski, B.; Sacher, O.; Magdziarz, T.; Bienfait, B. Development of a Battery of in Silico Prediction Tools for Drug-Induced Liver Injury from the Vantage Point of Translational Safety Assessment. Chem. Res. Toxicol. 2021, 34 (2), 601–615. 10.1021/acs.chemrestox.0c00423.

(17) Soldatow, V. Y.; Lecluyse, E. L.; Griffith, L. G.; Rusyn, I. In vitro Models for Liver Toxicity Testing. Toxicol. Res. (Camb). 2013, 2 (1), 23–39. 10.1039/c2tx20051a.

(18) Meng, Q. Three-Dimensional Culture of Hepatocytes for Prediction of Drug-Induced Hepatotoxicity. Expert Opin. Drug Metab. Toxicol. 2010, 6 (6), 733–746. 10.1517/17425251003674356.

(19) Sison-Young, R. L. C.; Mitsa, D.; Jenkins, R. E.; Mottram, D.; Alexandre, E.; Richert, L.; Aerts, H.; Weaver, R. J.; Jones, R. P.; Johann, E.; Hewitt, P. G.; Ingelman-Sundberg, M.; Goldring, C. E. P.; Kitteringham, N. R.; Park, B. K. Comparative Proteomic Characterization of 4 Human Liver-Derived Single Cell Culture Models Reveals Significant Variation in the Capacity for Drug Disposition, Bioactivation, and Detoxication. Toxicol. Sci. 2015, 147 (2), 412–424. 10.1093/toxsci/kfv136.

(20) Steger-Hartmann, T.; Raschke, M. Translating in Vitro to in Vivo and Animal to Human. Curr. Opin. Toxicol. 2020, 23–24, 6–10. 10.1016/j.cotox.2020.02.003.

(21) Kindrat, I.; Dreval, K.; Shpyleva, S.; Tryndyak, V.; de Conti, A.; Mudalige, T. K.; Chen, T.; Erstenyuk, A. M.; Beland, F. A.; Pogribny, I. P. Effect of Methapyrilene Hydrochloride on Hepatic Intracellular Iron Metabolism *in vivo* and *in vitro*. Toxicol. Lett. 2017, 281, 65–73. 10.1016/j.toxlet.2017.09.011.

(22) Graham, E. E.; Walsh, R. J.; Hirst, C. M.; Maggs, J. L.; Martin, S.; Wild, M. J.; Wilson, I. D.; Harding, J. R.; Kenna, J. G.; Peter, R. M.; Williams, D. P.; Park, B. K. Identification of the Thiophene Ring of Methapyrilene as a Novel Bioactivation-Dependent Hepatic Toxicophore. J. Pharmacol. Exp. Ther. 2008, 326 (2), 657–671. 10.1124/jpet.107.135483.

(23) Hamadeh, H. K.; Knight, B. L.; Haugen, A. C.; Sieber, S.; Amin, R. P.; Bushel, P. R.; Stoll, R.; Blanchard, K.; Jayadev, S.; Tennant, R. W.; Cunningham, M. L.; Afshari, C. A.; Paules, R. S. Methapyrilene Toxicity: Anchorage of Pathologic Observations to Gene Expression Alterations. Toxicol. Pathol. 2002, 30 (4), 470–482. 10.1080/01926230290105712.

(24) Mirsalis, J. C. Genotoxicity, Toxicity, and Carcinogenicity of the Antihistamine Methapyrilene (MTR 07216). Mutat. Res. Genet. Toxicol. 1987, 185 (3), 309–317. 10.1016/0165-1110(87)90022-4.

(25) Wright, P. S. R.; Briggs, K. A.; Thomas, R.; Smith, G. F.; Maglennon, G.; Mikulskis, P.; Chapman, M.; Greene, N.; Phillips, B. U.; Bender, A. Statistical Analysis of Preclinical Inter-Species Concordance of Histopathological Findings in the ETOX Database. Regul. Toxicol. Pharmacol. 2023, 138, 105308. 10.1016/j.yrtph.2022.105308.

(26) Olson, H.; Betton, G.; Robinson, D.; Thomas, K.; Monro, A.; Kolaja, G.; Lilly, P.; Sanders, J.; Sipes, G.; Bracken, W.; Dorato, M.; Van Deun, K.; Smith, P.; Berger, B.; Heller, A. Concordance of the Toxicity of Pharmaceuticals in Humans and in Animals. Regul. Toxicol. Pharmacol. 2000, 32 (1), 56–67. 10.1006/rtph.2000.1399.

(27) Fourches, D.; Barnes, J. C.; Day, N. C.; Bradley, P.; Reed, J. Z.; Tropsha, A. Cheminformatics Analysis of Assertions Mined from Literature That Describe Drug-Induced Liver Injury in Different Species. Chem. Res. Toxicol. 2010, 23 (1), 171–183. 10.1021/tx900326k.

(28) Bender, A.; Cortés-Ciriano, I. Artificial Intelligence in Drug Discovery: What Is Realistic, What Are Illusions? Part 1: Ways to Make an Impact, and Why We Are Not There Yet. Drug Discovery Today. Elsevier Ltd February 1, 2021, pp 511–524. 10.1016/j.drudis.2020.12.009.

(29) Van Norman, G. A. Limitations of Animal Studies for Predicting Toxicity in Clinical Trials: Is It Time to Rethink Our Current Approach? JACC Basic to Transl. Sci. 2019, 4 (7), 845–854. 10.1016/j.jacbts.2019.10.008.

(30) Cunningham, M. L. A Mouse Is Not a Rat Is Not a Human: Species Differences Exist. Toxicol. Sci. 2002, 70 (2), 157–158. 10.1093/toxsci/70.2.157.

(31) Thakkar, S.; Li, T.; Liu, Z.; Wu, L.; Roberts, R.; Tong, W. Drug-Induced Liver Injury Severity and Toxicity (DILIst): Binary Classification of 1279 Drugs by Human Hepatotoxicity. Drug Discov. Today 2020, 25 (1), 201–208. 10.1016/J.DRUDIS.2019.09.022.

(32) Chen, M.; Suzuki, A.; Thakkar, S.; Yu, K.; Hu, C.; Tong, W. DILIrank: The Largest Reference Drug List Ranked by the Risk for Developing Drug-Induced Liver Injury in Humans. Drug Discov. Today 2016, 21 (4), 648–653. 10.1016/J.DRUDIS.2016.02.015.

(33) Lawrence, E.; El-Shazly, A.; Seal, S.; Joshi, C. K.; Liò, P.; Singh, S.; Bender, A.; Sormanni, P.; Greenig, M. Understanding Biology in the Age of Artificial Intelligence. arXiv 2024, arXiv:2403.04106. 10.48550/arXiv.2403.04106

(34) Hewitt, M.; Enoch, S. J.; Madden, J. C.; Przybylak, K. R.; Cronin, M. T. D. Hepatotoxicity: A Scheme for Generating Chemical Categories for Read-across, Structural Alerts and Insights into Mechanism(s) of Action. Crit. Rev. Toxicol. 2013, 43 (7), 537–558. 10.3109/10408444.2013.811215.

(35) Ye, L.; Ngan, D. K.; Xu, T.; Liu, Z.; Zhao, J.; Sakamuru, S.; Zhang, L.; Zhao, T.; Xia, M.; Simeonov, A.; Huang, R. Prediction of Drug-Induced Liver Injury and Cardiotoxicity Using Chemical Structure and *in vitro* Assay Data. Toxicol. Appl. Pharmacol. 2022, 454. 10.1016/J.TAAP.2022.116250.

(36) Liu, A.; Walter, M.; Wright, P.; Bartosik, A.; Dolciami, D.; Elbasir, A.; Yang, H.; Bender, A. Prediction and Mechanistic Analysis of Drug-Induced Liver Injury (DILI) Based on Chemical Structure. Biol. Direct 2021, 16 (1), 1–15. 10.1186/s13062-020-00285-0.

(37) Mora, J. R.; Marrero-Ponce, Y.; García-Jacas, C. R.; Suarez Causado, A. Ensemble Models Based on QuBiLS-MAS Features and Shallow Learning for the Prediction of Drug-Induced Liver Toxicity: Improving Deep Learning and Traditional Approaches. Chem. Res. Toxicol. 2020, 33 (7), 1855–1873. 10.1021/ACS.CHEMRESTOX.0C00030/ASSET/IMAGES/LARGE/TX0C00030_0007.JPEG.

(38) Liu, A.; Seal, S.; Yang, H.; Bender, A. Using Chemical and Biological Data to Predict Drug Toxicity. SLAS Discov. 2023. 10.1016/J.SLASD.2022.12.003.

(39) Seal, S.; Yang, H.; Trapotsi, M.-A.; Singh, S.; Carreras-Puigvert, J.; Spjuth, O.; Bender, A. Merging Bioactivity Predictions from Cell Morphology and Chemical Fingerprint Models Using Similarity to Training Data. J. Cheminform. 2023, 15 (1), 56. 10.1186/s13321-023-00723-x.

(40) Kaito, S.; Takeshita, J.; Iwata, M.; Sasaki, T.; Hosaka, T.; Shizu, R.; Yoshinari, K. Utility of Human Cytochrome P450 Inhibition Data in the Assessment of Drug-Induced Liver Injury. Xenobiotica 2024, 1–30. 10.1080/00498254.2024.2312505.

(41) Rao, M.; Nassiri, V.; Alhambra, C.; Snoeys, J.; Van Goethem, F.; Irrechukwu, O.; Aleo, M. D.; Geys, H.; Mitra, K.; Will, Y. AI/ML Models to Predict the Severity of Drug-Induced Liver Injury for Small Molecules. Chem. Res. Toxicol. 2023, 36, 1129–1139. 10.1021/acs.chemrestox.3c00098.

(42) Aleo, M. D.; Shah, F.; Allen, S.; Barton, H. A.; Costales, C.; Lazzaro, S.; Leung, L.; Nilson, A.; Obach, R. S.; Rodrigues, A. D.; Will, Y. Moving beyond Binary Predictions of Human Drug-Induced Liver Injury (DILI) toward Contrasting Relative Risk Potential. Chem. Res. Toxicol. 2020, 33 (1), 223–238. 10.1021/acs.chemrestox.9b00262.

(43) Chavan, S.; Scherbak, N.; Engwall, M.; Repsilber, D. Predicting Chemical-Induced Liver Toxicity Using High-Content Imaging Phenotypes and Chemical Descriptors: A Random Forest Approach. Chem. Res. Toxicol. 2020, 33 (9), 2261–2275. 10.1021/ACS.CHEMRESTOX.9B00459/SUPPL_FILE/TX9B00459_SI_003.TXT.

(44) Seal, S.; Carreras-Puigvert, J.; Trapotsi, M. A.; Yang, H.; Spjuth, O.; Bender, A. Integrating Cell Morphology with Gene Expression and Chemical Structure to Aid Mitochondrial Toxicity Detection. Commun. Biol. 2022, 5 (1), 858. 10.1038/s42003-022-03763-5.

(45) Seal, S.; Yang, H.; Vollmers, L.; Bender, A. Comparison of Cellular Morphological Descriptors and Molecular Fingerprints for the Prediction of Cytotoxicity- And Proliferation-Related Assays. Chem. Res. Toxicol. 2021, 34 (2), 422–437. 10.1021/acs.chemrestox.0c00303.

(46) Seal, S.; Spjuth, O.; Hosseini-Gerami, L.; García-Ortegón, M.; Singh, S.; Bender, A.; Carpenter, A. E. Insights into Drug Cardiotoxicity from Biological and Chemical Data: The First Public Classifiers for FDA Drug-Induced Cardiotoxicity Rank. J. Chem. Inf. Model. 2024. 10.1021/ACS.JCIM.3C01834.

(47) Mulliner, D.; Schmidt, F.; Stolte, M.; Spirkl, H. P.; Czich, A.; Amberg, A. Computational Models for Human and Animal Hepatotoxicity with a Global Application Scope. Chem. Res. Toxicol. 2016, 29 (5), 757–767. 10.1021/ACS.CHEMRESTOX.5B00465

(48) Liu, J.; Mansouri, K.; Judson, R. S.; Martin, M. T.; Hong, H.; Chen, M.; Xu, X.; Thomas, R. S.; Shah, I. Predicting Hepatotoxicity Using ToxCast *in vitro* Bioactivity and Chemical Structure. Chem. Res. Toxicol. 2015, 28 (4), 738–751. 10.1021/TX500501H.

(49) Ambe, K.; Ishihara, K.; Ochibe, T.; Ohya, K.; Tamura, S.; Inoue, K.; Yoshida, M.; Tohkin, M. In Silico Prediction of Chemical-Induced Hepatocellular Hypertrophy Using Molecular Descriptors. Toxicol. Sci. 2018, 162 (2), 667–675. 10.1093/TOXSCI/KFX287.

(50) He, S.; Ye, T.; Wang, R.; Zhang, C.; Zhang, X.; Sun, G.; Sun, X. An In Silico Model for Predicting Drug-Induced Hepatotoxicity. Int. J. Mol. Sci. 2019, 20 (8). 10.3390/IJMS20081897.

(51) Hemmerich, J.; Troger, F.; Füzi, B.; F.Ecker, G. Using Machine Learning Methods and Structural Alerts for Prediction of Mitochondrial Toxicity. Mol. Inform. 2020, 39 (5). 10.1002/minf.202000005.

(52) McLoughlin, K. S.; Jeong, C. G.; Sweitzer, T. D.; Minnich, A. J.; Tse, M. J.; Bennion, B. J.; Allen, J. E.; Calad-Thomson, S.; Rush, T. S.; Brase, J. M. Machine Learning Models to Predict Inhibition of the Bile Salt Export Pump. J. Chem. Inf. Model. 2021, 61 (2), 587–602. 10.1021/ACS.JCIM.0C00950.

(53) Mazzolari, A.; Vistoli, G.; Testa, B.; Pedretti, A. Prediction of the Formation of Reactive Metabolites by A Novel Classifier Approach Based on Enrichment Factor Optimization (EFO) as Implemented in the VEGA Program. Molecules 2018, 23 (11). 10.3390/MOLECULES23112955.

(54) Zhang, C.; Zhao, Y.; Yu, M.; Qin, J.; Ye, B.; Wang, Q. Mitochondrial Dysfunction and Chronic Liver Disease. Curr. Issues Mol. Biol. 2022, 44 (7), 3156–3165. 10.3390/cimb44070218.

(55) Kenna, J. G.; Taskar, K. S.; Battista, C.; Bourdet, D. L.; Brouwer, K. L. R.; Brouwer, K. R.; Dai, D.; Funk, C.; Hafey, M. J.; Lai, Y.; Maher, J.; Pak, Y. A.; Pedersen, J. M.; Polli, J. W.; Rodrigues, A. D.; Watkins, P. B.; Yang, K.; Yucha, R. W. Can Bile Salt Export Pump Inhibition Testing in Drug Discovery and Development Reduce Liver Injury Risk? An International Transporter Consortium Perspective. Clin. Pharmacol. Ther. 2018, 104 (5), 916–932. 10.1002/cpt.1222.

(56) Attia, S. M. Deleterious Effects of Reactive Metabolites. Oxid. Med. Cell. Longev. 2010, 3 (4), 238–253. 10.4161/oxim.3.4.13246.

(57) Seal, S.; Trapotsi, M.-A.; Subramanian, V.; Spjuth, O.; Greene, N.; Bender, A.-D. PKSmart: An Open-Source Computational Model to Predict in Vivo Pharmacokinetics of Small Molecules. bioRxiv 2024, 2024.02.02.578658. 10.1101/2024.02.02.578658.

(58) Varnek, A.; Gaudin, C.; Marcou, G.; Baskin, I.; Pandey, A. K.; Tetko, I. V. Inductive Transfer of Knowledge: Application of Multi-Task Learning and Feature Net Approaches to Model Tissue-Air Partition Coefficients. J. Chem. Inf. Model. 2009, 49 (1), 133–144. 10.1021/CI8002914/SUPPL_FILE/CI8002914_SI_001.PDF.

(59) Martin, M. T. and R. Judson. U.S. Environmental Protection Agency, Washington, DC, 2010. Release User-Friendly Web-Based Tool for Mining ToxRefDB.

(60) Sakuratani, Y.; Zhang, H. Q.; Nishikawa, S.; Yamazaki, K.; Yamada, T.; Yamada, J.; Gerova, K.; Chankov, G.; Mekenyan, O.; Hayashi, M. Hazard Evaluation Support System (HESS) for Predicting Repeated Dose Toxicity Using Toxicological Categories. SAR QSAR Environ. Res. 2013, 24 (5), 351–363. 10.1080/1062936X.2013.773375.

(61) Williams, D. P.; Lazic, S. E.; Foster, A. J.; Semenova, E.; Morgan, P. Predicting Drug-Induced Liver Injury with Bayesian Machine Learning. Chem. Res. Toxicol. 2020, 33 (1), 239–248. 10.1021/acs.chemrestox.9b00264.

(62) Horne, R.I.; Wilson-Godber, J.; Díaz, A. G.; Brotzakis, Z. F.; Seal, S.; Gregory, R. C.; Possenti, A.; Chia, S.; Vendruscolo, M. Using Generative Modeling to Endow with Potency Initially Inert Compounds with Good Bioavailability and Low Toxicity. J. Chem. Inf. Model. 2024 (in press). 10.1021/acs.jcim.3c01777.

(63) Smit, I. A.; Afzal, A. M.; Allen, C. H. G.; Svensson, F.; Hanser, T.; Bender, A. Systematic Analysis of Protein Targets Associated with Adverse Events of Drugs from Clinical Trials and Postmarketing Reports. Chem. Res. Toxicol. 2021, 34 (2), 365–384. 10.1021/ACS.CHEMRESTOX.0C00294/ASSET/IMAGES/LARGE/TX0C00294_0009.JPEG.

(64) RDKit (v2022.09.5) https://www.rdkit.org/ (accessed Apr 18, 2023).

(65) Ropp, P. J.; Kaminsky, J. C.; Yablonski, S.; Durrant, J. D. Dimorphite-DL: An Open-Source Program for Enumerating the Ionization States of Drug-like Small Molecules. J. Cheminform. 2019, 11 (1), 1–8. 10.1186/s13321-019-0336-9.

(66) scikit-learn: machine learning in Python — scikit-learn 1.2.0 documentation https://scikit-learn.org/stable/index.html (accessed Jan 3, 2023).

(67) Rogers, D.; Hahn, M. Extended-Connectivity Fingerprints. J. Chem. Inf. Model. 2010, 50 (5), 742–754. 10.1021/ci100050t.

(68) Durant, J. L.; Leland, B. A.; Henry, D. R.; Nourse, J. G. Reoptimization of MDL Keys for Use in Drug Discovery. J. Chem. Inf. Comput. Sci. 2002, 42 (6), 1273–1280. 10.1021/ci010132r.

(69) Moriwaki, H.; Tian, Y. S.; Kawashita, N.; Takagi, T. Mordred: A Molecular Descriptor Calculator. J. Cheminform. 2018, 10 (1), 1–14. 10.1186/s13321-018-0258-y.

(70) The DeepChem Project — deepchem 2.7.2.dev documentation https://deepchem.readthedocs.io/en/latest/ (accessed Mar 28, 2024).

(71) Ramsudar, B.; Eastman, P.; Walters, P.; Pande, V. Deep Learning for Life Sciences. 2019, 222.

(72) Fierz, W.; Bossuyt, X. Likelihood Ratio Approach and Clinical Interpretation of Laboratory Tests. Front. Immunol. 2021, 12, 1170. 10.3389/fimmu.2021.655262.

(73) SHAP. Welcome to the SHAP documentation — SHAP latest documentation https://shap.readthedocs.io/en/latest/index.html (accessed May 17, 2023).

(74) A.; Chang, P.; Devuni, D.; Bichoupan, K.; Kesar, V.; Branch, A. D.; Oh, W. K.; D. Galsky, M.; Ahmad, J.; A. Odin, J. Real World Experience of Drug Induced Liver Injury in Patients Undergoing Chemotherapy. J. Clin. Gastroenterol. Hepatol. 2018, 02 (03). 10.21767/2575-7733.1000047.

(75) Chen, M.; Borlak, J.; Tong, W. A Model to Predict Severity of Drug-Induced Liver Injury in Humans. Hepatology 2016, 64 (3), 931–940. 10.1002/hep.28678.

(76) Khalili, H.; Soudbakhsh, A.; Talasaz, A. H. Severe Hepatotoxicity and Probable Hepatorenal Syndrome Associated with Sulfadiazine. Am. J. Heal. Pharm. 2011, 68 (10), 888–892. 10.2146/ajhp100516.

(77) Galletti, E.; Magnani, M.; Renzulli, M. L.; Botta, M. Paclitaxel and Docetaxel Resistance: Molecular Mechanisms and Development of New Generation Taxanes. ChemMedChem 2007, 2 (7), 920–942. 10.1002/cmdc.200600308.

(78) Badwan, B. A.; Liaropoulos, G.; Kyrodimos, E.; Skaltsas, D.; Tsirigos, A.; Gorgoulis, V. G. Machine Learning Approaches to Predict Drug Efficacy and Toxicity in Oncology. Cell Reports Methods 2023, 3 (2). 10.1016/j.crmeth.2023.100413.

(79) Barelier, S.; Eidam, O.; Fish, I.; Hollander, J.; Figaroa, F.; Nachane, R.; Irwin, J. J.; Shoichet, B. K.; Siegal, G. Increasing Chemical Space Coverage by Combining Empirical and Computational Fragment Screens. ACS Chem. Biol. 2014, 9 (7), 1528–1535. 10.1021/cb5001636.

(80) HESI Insights - February 2023 - HESI - Health and Environmental Sciences Institute https://hesiglobal.org/hesi-insights-february-2023/ (accessed May 29, 2023).

(81) Garcia-Ortegon, M.; Singh, S.; Bender, A.; Bacallado, S. Calibrated Prediction of Scarce Adverse Drug Reaction Labels with Conditional Neural Processes Garcia-Ortegon et Al.

(82) Plowright, A. T.; Johnstone, C.; Kihlberg, J.; Pettersson, J.; Robb, G.; Thompson, R. A. Hypothesis Driven Drug Design: Improving Quality and Effectiveness of the Design-Make-Test-Analyse Cycle. Drug Discov. Today 2012, 17 (1–2), 56–62. 10.1016/j.drudis.2011.09.012.

