## Supplementary Figures for "Improved Detection of Drug-Induced Liver Injury by Integrating Predicted *in vivo* and *in vitro* Data"

Machine Learning, Toxicity, DILI, *in vivo*, *in vitro*, Drug-Induced Liver Injury

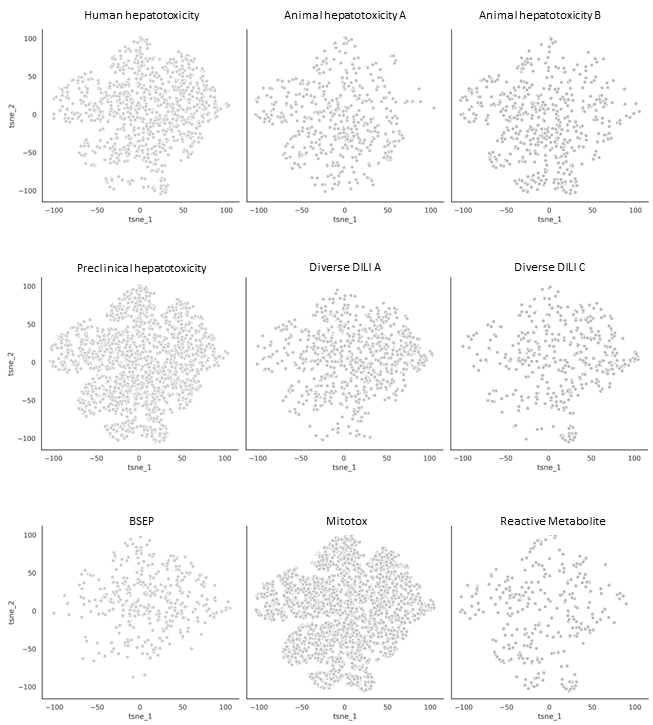

Figure S1: Distribution of 12,133 compounds in proxy-DILI dataset in each of the 9 labels as represented by the t-SNE plot of physicochemical space.

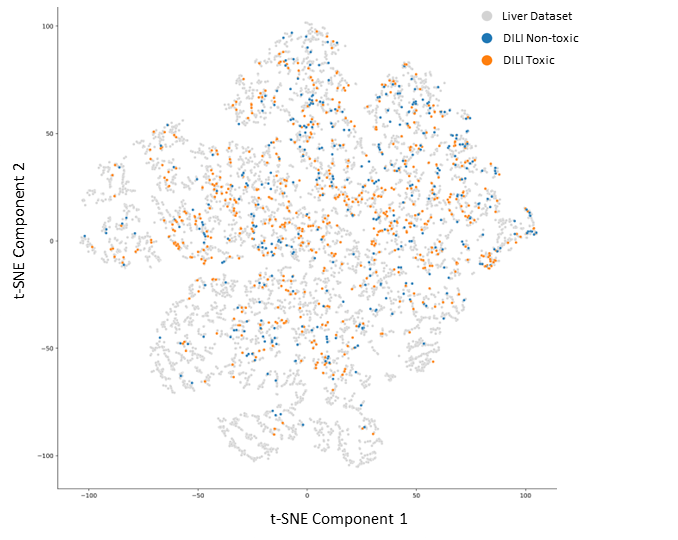

Figure S2: Distribution of 12,133 compounds in proxy-DILI dataset and 1,111 in the gold standard DILI dataset as represented by the t-SNE plot of physicochemical space. Overall, DILI compounds were representative of the physicochemical space of the compounds in the proxy-DILI dataset.

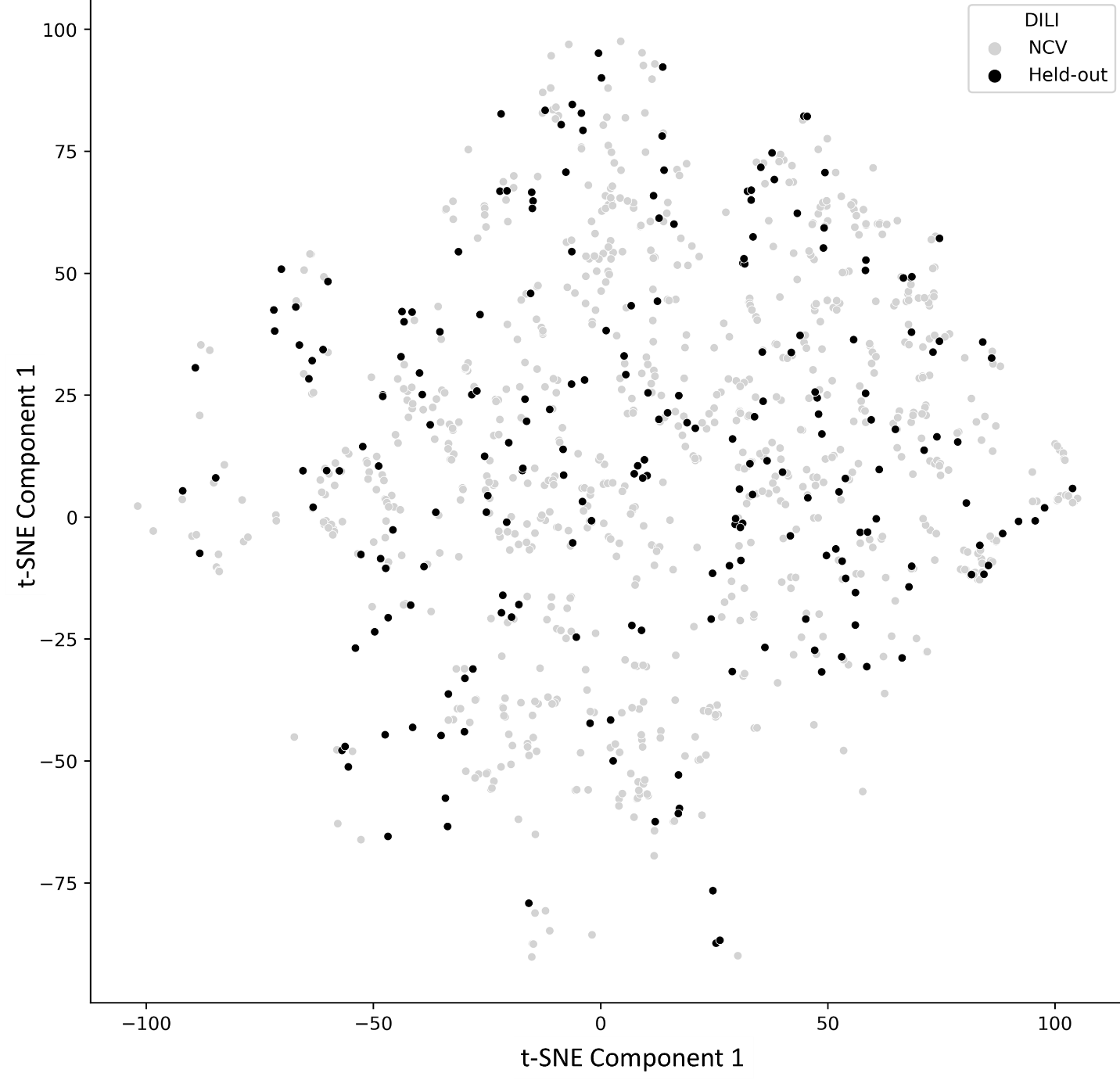

Figure S3: Distribution of the held-out DILI test set (223 compounds) and the training DILI data (888 compounds) as represented by the t-SNE plot of physicochemical space. Overall, compounds in the held-out DILI test set were representative of the physicochemical space of the compounds in the training DILI data dataset.

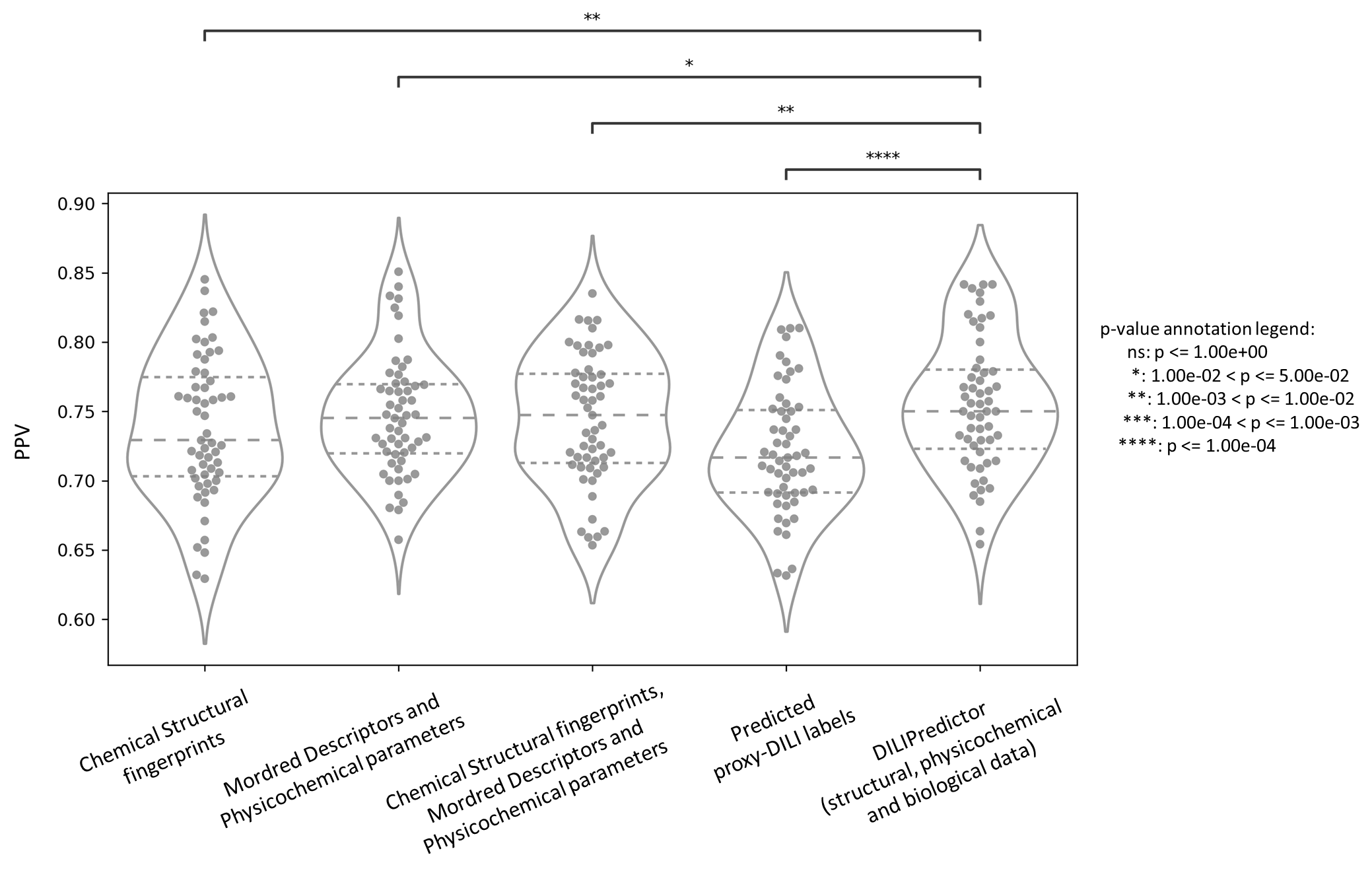

Figure S4: Performance metric positive predictive value for combination models from 55 held-out test sets from repeated nested cross-validation using (a) selected 189-bit structural fingerprints, (b) selected 668 molecular descriptors, (c) selected 189-bit structural fingerprints and selected 668 molecular descriptors, (d) predicted nine proxy-DILI labels and 2 PK parameters, and (e) a combination of all three features spaces, compared with a paired t-test.

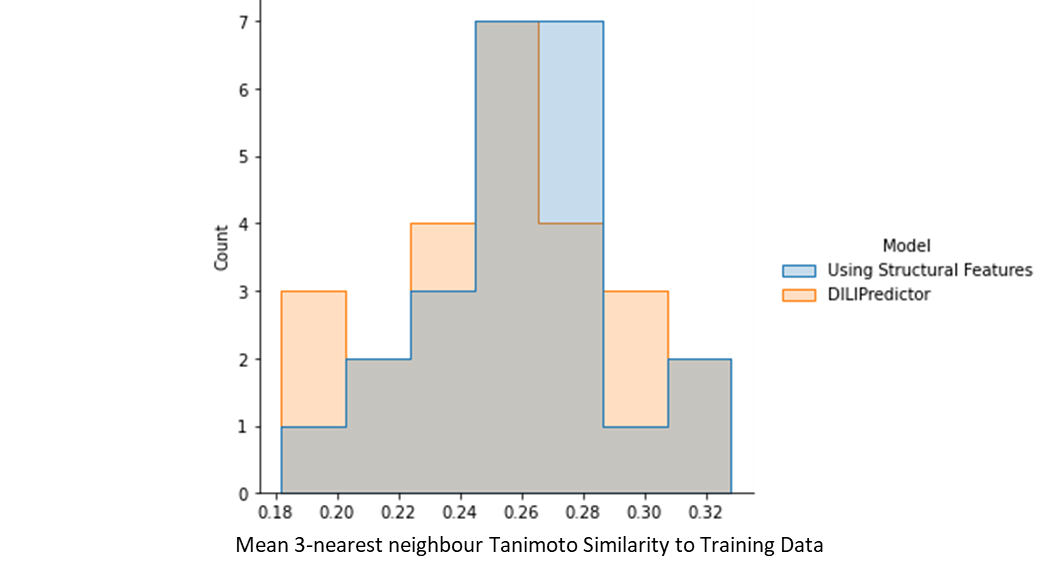

Figure S5: Distribution of 3-nearsed neighbour Tanimoto similarity for toxic compounds detected correctly in the early stage (from the top 29 compounds) with a low false positive rate for models using only structural features compared to DILIPredictor models using all feature spaces.

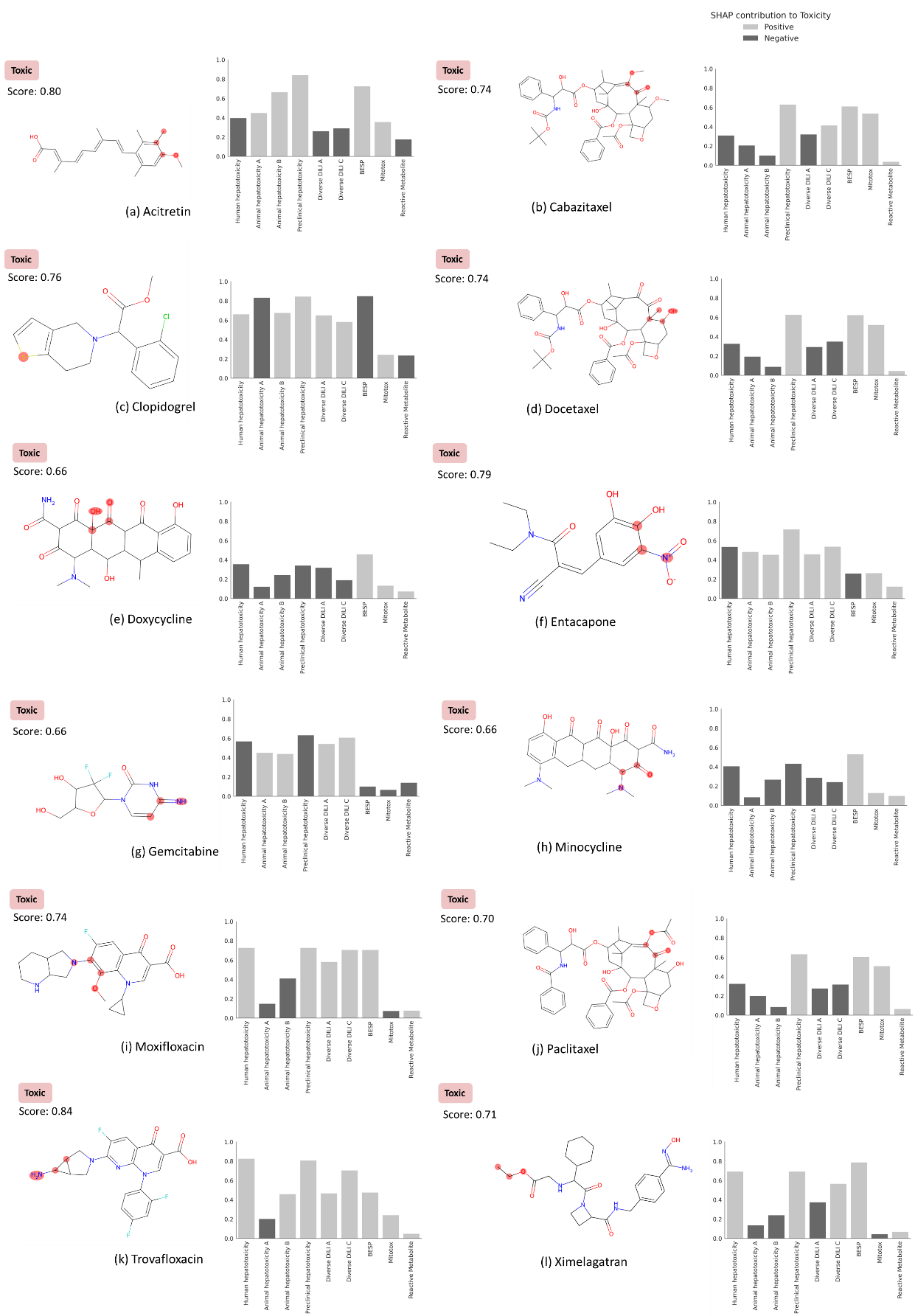

Figure S6: MACCS substructure positively contributing to DILI (highlighted) and the proxy-DILI predictions that are positively and negatively for contributing to a prediction (SHAP values) when using DILIPredictor for 12 compounds known to cause DILI in humans (in training dataset). The highest positive contribution from the MACCS substructure is highlighted with the chemical structure. See Figure 7 for two compounds known to cause DILI and two compounds which do not cause DILI in humans (from external test set).
